# High-throughput *in vivo* synaptic connectivity mapping of neuronal micro-circuits using two-photon holographic optogenetics and compressive sensing

**DOI:** 10.1101/2023.09.11.557026

**Authors:** I-Wen Chen, Chung Yuen Chan, Phillip Navarro, Vincent de Sars, Emiliano Ronzitti, Karim Oweiss, Dimitrii Tanese, Valentina Emiliani

## Abstract

Understanding the intricate synaptic connectivity in living neural circuits is crucial for unraveling the relationship between network structure and function, as well as its evolution during development, learning, and recovery from injury. However, current methodologies for identifying connected neurons *in vivo* suffer from limitations, particularly with regards to their throughput. In this study, we introduce a groundbreaking framework for *in vivo* connectivity mapping that combines two-photon holographic optogenetics for activating single or multiple potential presynaptic neurons, whole-cell recording of postsynaptic responses, and a compressive sensing strategy for efficiently retrieving individual postsynaptic neurons’ responses when multiple potential presynaptic neurons are simultaneously activated. The approach was validated in the layer 2/3 of the visual cortex in anesthetized mice, enabling rapid probing of up to 100 cells in approximately 5 minutes. By identifying tens of synaptic pairs, including their connection strength, kinetics, and spatial distribution, this method showcases its potential to significantly advance circuit reconstruction in large neuronal networks with minimal invasiveness. Moreover, through simultaneous multi-cell stimulation and compressive sensing, we demonstrate up to a three-fold reduction in the number of required measurements to infer connectivity with limited loss in accuracy, thereby enabling high-throughput connectivity mapping *in vivo*. These results pave the way for a more efficient and rapid investigation of neuronal circuits, leading to deeper insights into brain function and plasticity.

## Introduction

The elucidation of synaptic connectivity among neurons is of utmost importance for comprehending the neural circuitry responsible for mediating behavior, driving extensive efforts towards the development and enhancement of experimental approaches for neuronal circuit connectivity mapping, both *in vitro* and *in vivo* ^1–3^. *In vivo* investigations under normal physiological conditions is essential to avoid disruption of circuit function when connections are severed during tissue slicing procedures postmortem ^4–6^. Most importantly, mapping connections in a defined cell population *in vivo* repeatedly enables tracking the changes in connectivity over time, thereby correlating synaptic plasticity with measurable changes in behavior, such as during development, learning, memory formation or recovery from injury.

Although correlated activity in neuronal subpopulations from fluorescence imaging or extracellular multiple single-unit recordings ^7–9^ provides coarse and indirect information about connectivity, they do not provide information of individual monosynaptic connections happening at very short latencies (typically < 5 ms) ^10–12^. Ascertaining its presence can only be achieved using causal induction of action potentials (AP) at the presynaptic level and intracellular recording of postsynaptic response (current or voltage) ^13–20^. However, this approach has major limitations. First, it precludes the ability to examine connectivity between the cells in future experiments. Second, it is difficult to achieve and maintain either single or multiple intracellular recordings for prolonged periods, thereby lowering the yield of the experiment. As such, this approach has so far been limited to investigating connections among a maximum of 12 neurons *in vitro* ^17^ and up to 4 neurons per experiment *in vivo* ^20^.

Attempts to increase the number of probed pairs have used optical approaches such as glutamate uncaging ^21^ and optogenetics ^22^. In the first case, presynaptic spiking was induced using single-photon (1P) ^23–28^ or two-photon (2P) ^29–31^ multi-point photolysis of caged glutamate around dendrites or soma. Under 2P excitation, glutamate uncaging enabled AP induction at near cellular resolution and consequently probing connectivity over several hundreds of potential presynaptic cells ^30,31^. However, glutamate uncaging lacks control over the timing and number of induced spikes as well as the diffusion of the caged compound, which can affect the spatial resolution and cell-type specificity of presynaptic activation. Moreover, the need for perfusing caged glutamate hinders the possibility to use these approaches *in vivo*.

Alternatively, presynaptic activity can be induced *in vitro* using 1P optogenetic stimulation of dendrites, soma, or axons ^32–36^. Leveraging precise control of the channel kinetics and light-sensitivity of opsins, 1P optogenetic stimulation has achieved optical control of the timing and number of induced presynaptic spikes ^34,37^, but cellular resolution was achievable only at shallow depths and under sparse opsin-expression, limiting the study of connectivity *in vivo* to a single presynaptic neuron ^32,33^.

In contrast, 2P optogenetics enables both *in vitro* and *in vivo* AP generation at cellular resolution ^38^ and can be a powerful tool to precisely manipulate larger ensembles of potential presynaptic neurons when probing synaptic connectivity. Specifically, 2P illumination methods using laser scanning ^39–47^ or parallel light-patterning ^48–64^ ensures scattering-resistant actuation of opsins deep in brain tissue, which can reach cellular resolution when combined with soma-restricted opsins ^55,58,65,66^. Moreover, 2P stimulation of opsins with high light-sensitivity and fast channel kinetics ensures temporally precise AP generation of millisecond latency and submillisecond jitter *in vitro* ^54,55^ and *in vivo* ^42,61,67^, which is of key importance to correlate the timing of presynaptic stimulation with postsynaptic responses.

Initial proof-of-concept experiments have allowed for the establishment of optogenetic connectivity mapping ^55,58,68^, and have been followed by recent studies which have used this approach to elucidate the synaptic organization of the medial prefrontal cortex in mice ^69^, to characterize intralaminar and translaminar monosynaptic connections in the mouse visual cortex ^70^, and to investigate fast undulatory locomotion in zebrafish ^71^. With the exception of the last study performed on transparent zebrafish larvae *in vivo*, all these works have been carried out *in vitro.* Doing so in the mammalian brain *in vivo* has some major challenges to overcome such as the fast time-varying neuromodulatory activity, whole-cell patching instability, and membrane potential fluctuation of spontaneous activity during UP and DOWN states. These factors, among others, largely affect the probability and temporal properties of presynaptic APs, thus preventing accurate measurements of the evoked postsynaptic responses, which are typically of small amplitudes of ∼1 mV or a few picoamperes. These limitations increase the number of required trials for noise reduction through trial averaging, require long recording durations that are hardly compatible with whole-cell patch-clamp, and make *in vivo* large-scale mapping of synaptic connectivity out of reach.

Here, we present a novel experimental and computational strategy for *in vivo* high-throughput connectivity mapping where we combine optogenetics and temporally-focused 2D and 3D holographic light-multiplexing ^48^ to control the spiking activity of a single or multiple targets with single-cell resolution, millisecond resolution, and sub-millisecond precision. We also demonstrate that the speed and throughput of the method can be increased by combining 2P optogenetic multi-cell stimulation for presynaptic activation with compressive sensing (CS) ^36,72–75^—an approach that uses sparsity and incoherent sampling to find a solution to an underdetermined linear system ^76,77^.

We first demonstrate that mapping can be achieved using rapid, sequential stimulation of single potential presynaptic neurons expressing the fast, light-sensitive soma-restricted opsin ST-ChroME ^66^ and whole-cell recording for postsynaptic readout. The approach allows to estimate connection rates and synaptic properties of L2/3 intralaminar connectivity, probing up to 100 potential presynaptic neurons per experiment and identifying several tens of connected pairs, their synaptic strength and spatial distribution. Leveraging the innate scarcity of cortical neuronal connections, we then demonstrate that CS improves the efficiency and recording time by showing that identification of synaptic pairs is feasible using only a third of the measurements needed using single-cell sequential activation, with more than half of the connected pairs retrieved (recall≈60%) at an average false positive below 40%. To our knowledge, our results represent the first *in vivo* demonstration of high-throughput connectivity mapping at cellular resolution in mammalian brain.

## Results

### The optical system

Experiments were carried out in a custom-built optical system (Fig. 1A) which included a path for 2P galvanometric scan imaging and a path for 2P holographic stimulation (see Methods). The holographic system enabled generating single or multiple temporally-focused circular spots by using computer-generated holography via a two-step phase modulation process ^48,56^. First, the wavefront of the stimulation laser beam was modulated by a static phase mask to generate a circle-shaped holographic spot (12 µm in diameter, Fig. 1B) on a diffraction grating to implement temporal focusing and improve the axial resolution ^78,79^ (Fig. 1B). Second, the temporally focused holographic light-spot was multiplexed at multiple positions by a liquid crystal spatial light modulator (SLM). As for light source, we utilized a low-repetition rate (500 kHz), high-power fiber amplifier laser. The maximum power of stimulation laser was 10-50 W at the source and ∼1-5 W at the exit of the microscope objective, which allowed the generation of several tens of holographic spots within a field of view (FOV) of 350 µm x 350 µm x 100 µm.

**Figure 1:**
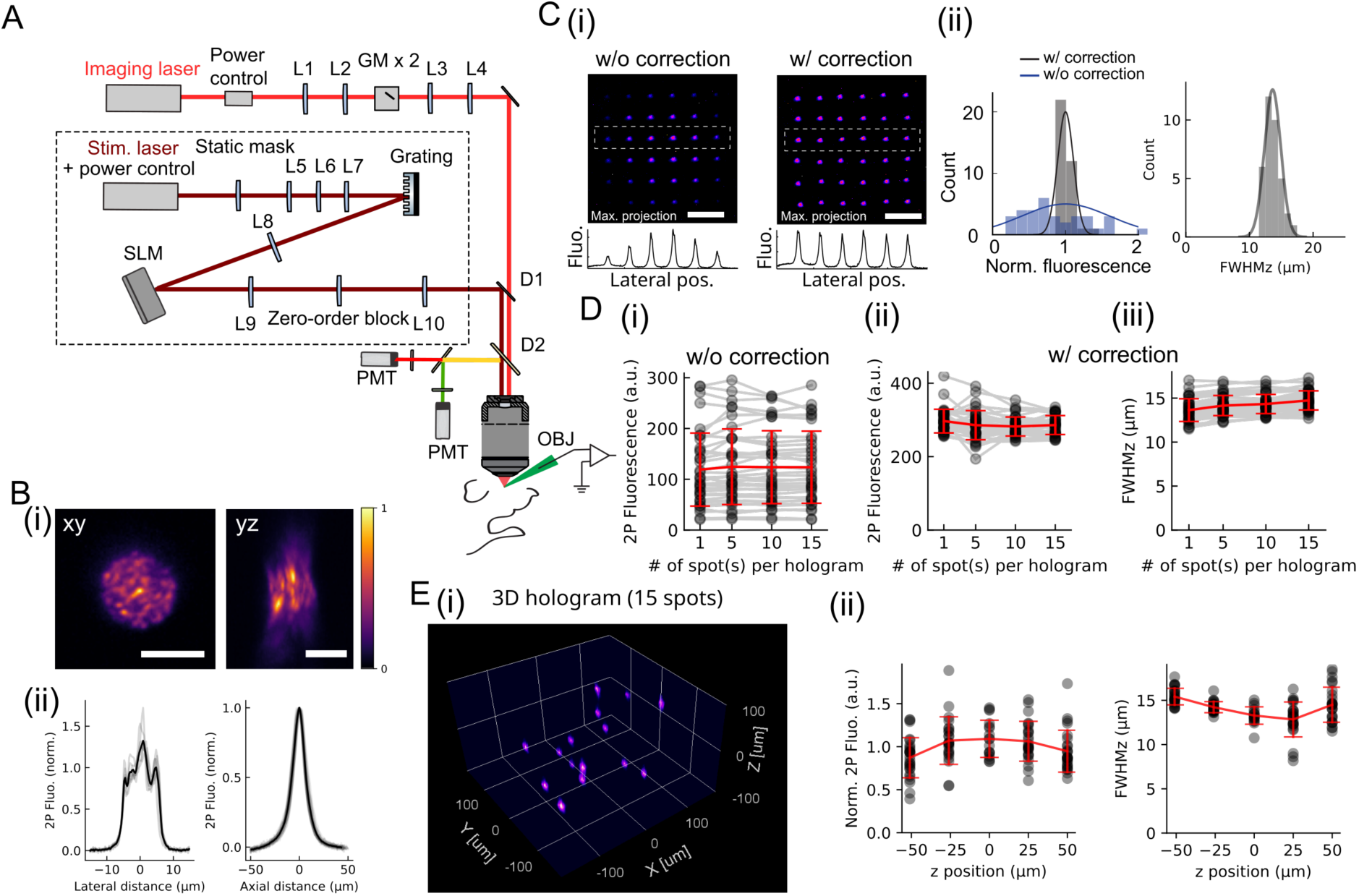
Optical system and characterization of holographic spots. (A) Schematic of the optical system comprising a path for 2P scan imaging (red light-beam) and a path for 2P holographic stimulation (dark red light-beam). L: lens, GM: galvanometric mirrors, D: dichroic mirror, PMT: photomultiplier tube, SLM: spatial light modulator, OBJ: objective lens. For the two paths we used respectively a Ti:sapphire tunable laser or a fixed wavelength fiber amplified laser at 1030 nm. (B) (i): Top and side view of the two-photon excited fluorescence generated by a holographic spot of 12 μm diameter on a thin rhodamine layer. (ii): Corresponding normalized lateral (12 μm FWHM) and axial profile (15 μm FWHM). Scale bar: 10 μm. Individual lateral and axial profiles from 7 and 36 holographic spots, respectively, denoted by light grey lines. Average profiles in black. (C) A sequence of 36 holograms were calculated to generate 36 single spots distributed over the FOV (300 μm x 300 μm) without and with compensation of the position-dependent diffraction efficiency. (i): Max. projection image of the 36 light-spots and intensity profile across the marked ROIs without and with compensation. Scale bar: 100 μm. (ii): Left: Distribution of the normalized integrated fluorescence spot intensities generated by the sequence of holographic spots with (grey) and without (blue) diffraction efficiency compensation and corresponding gaussian fits (standard deviation of 10.9% and 60.4% with and w/o correction respectively). Right: Distribution of the axial resolution (FWHM of the axial profile) of the holographic spots’ fluorescence. (D) Sequences of holograms were calculated to generate 1, 5, 10 or 15 spot(s) distributed over the FOV (300 μm x 300 μm) (i,ii): Fluorescence spot intensities as a function of number of spots per hologram, without (i) and with (ii) correction of diffraction efficiency; (iii): Holographic spots axial resolution as a function of number of spots per hologram. Fluorescence intensity values were extracted from n=36, 40, 20 and 15 holograms each generating 1,5,10,15 spot(s) respectively. Axial resolution values were extracted from n=36, 20, 10 and 8 holograms each generating 1,5,10,15 spot(s) respectively. Grey dots represent individual lateral spot position within the FOV, average values are in red (error bars: mean±s.d). (E) Multi-spot patterns of temporally-focused holographic spots distributed in 3 dimensions. (i): Example image of a 3D hologram with 15 spots in a 3D volume of 300 μm x 300 μm x 100 μm. (ii): 2P fluorescence and optical axial resolution at different axial positions (mean±s.d.). Points are extracted from 10 holograms, each generating 15 spots.

To guarantee the homogeneities of spots, both in size and intensities over the full FOV, we compensated the diffraction efficiency losses caused by the SLM by adjusting the total laser power and relative power ratio between targets, following procedures previously described ^55,56^ (see Methods). After this correction, the variations in fluorescence intensity across the FOV were limited to 9-14% (s.d.) of the mean fluorescence for single-spot and multi-spot generation (p=0.29 for fluorescence intensity, one-way ANOVA for spot number of 1, 5, 10, and 15, n=36 spot positions from 36, 40, 20 and 15 holograms, respectively) (Fig. 1C, D and Fig. S1A-F). Similarly, the spot shapes were homogenous over the 3D FOV and the axial profiles showed a FWHM ≈ 15 µm, not majorly affected by lateral and axial position (Fig. 1C, E and Fig. S1G-I) and by the number of spots generated (Fig. 1D, p=0.25, one-way ANOVA for spot number of 1, 5, 10, and 15, n=36 spot positions from 36, 20, 10 and 8 holograms, respectively) (FWHM 1-spot: 13.6 ± 1.3 µm, 10-spot: 14.3 ± 1.1 µm, mean ± s.d.).

### AP properties upon single-cell and multi-cell stimulation

To examine the stimulation conditions enabling reliable AP generation in a fast and temporally precise manner, we measured the AP properties upon shining 2P holographic spots onto single or multiple neurons expressing the fast and light-sensitive opsin ST-ChroME of soma-restricted expression ^66^. We expressed the opsin by using viral injection in layer 2/3 (L2/3) of mouse V1 (see Methods). By using 2P-guided whole-cell or cell-attached recording, we measured the spiking activity upon 2P holographic stimulation of opsin-positive neurons (see Methods), identified from the red mRuby fluorescence and monitored the main electrophysiological properties of the patched neurons (Table 1, Fig. S2A). These features, in particular resting membrane potential and spontaneous spiking rate, were used to recognize putative pyramidal cells among the patched neurons and restrict our characterization to this cell type. Of note, compared to opsin-negative ones, opsin-expressing neurons showed an increased AP threshold and input resistance (p<0.05, two-sample t-test for opsin-negative and opsin-positive cells, Table 1) and the latter may indicate an enhanced neuronal excitability after prolonged cross-talk activation of opsins induced by the scanning of the imaging laser for 2P-guided patching or identifying opsin-positive cells ^80^.

**Table 1:** Electrophysiological properties of opsin-negative and opsin-positive neurons.

|  | Opsin-negative cells | Opsin-positive cells | p-value of 2-sample t-test |
| --- | --- | --- | --- |
| Average membrane potential (mV) | -74.5±1.4 (n=9) | -74.5±0.8 (n=12) | 0.99 |
| Resting membrane potential (mV) | -74.5±1.4 (n=9) | -74.5±0.8 (n=12) | 0.99 |
| Input resistance (MΩ) | 64.4±5.8 (n=8) | 98.2±7.4 (n=12) | 0.0041 |
| Membrane time constant (ms) | 5.1±0.7 (n=8) | 5.0±0.7 (n=12) | 0.95 |
| Membrane capacitance (pF) | 78.8±8.3 (n=8) | 52.7±5.1 (n=12) | 0.011 |
| Rheobase (pA) | 268.8±21.0 (n=8) | 286.1±23.8 (n=12) | 0.61 |
| AP threshold (mV) | -37.3±2.0 (n=8) | -25.3±2.4 (n=12) | 0.0024 |
| AP peak amplitude (mV) | 67.3±3.8 (n=8) | 50.8±2.7 (n=12) | 0.0018 |
| AP half-width (ms) | 1.4±0.06 (n=8) | 1.5±0.1 (n=12) | 0.28 |
| Spontaneous spiking rate (Hz) | 0.00±0.00 (n=9) | 0.03±0.03 (n=12) | 0.40 |
| Evoked spiking rate (Hz) | 31.3±2.8 (n=6) | 33.6±3.6 (n=10) | 0.67 |

Data were presented as mean±s.e.m.; 9 opsin-negative cells and 12 opsin-positive cells.
Membrane potential was corrected for a liquid junction potential of 11.9 mV.
Evoked spiking rates were accessed as the spiking rates upon current injection of rheobase+100 pA.

We characterized photoinduced spiking activity using 3 holographic patterns targeting either the patched opsin-expressing neuron only (hereafter indicated as pattern ‘1 target’), or the somata of 10 surrounding neurons (pattern ‘10 targets’), or both (pattern ‘10+1 targets’) (Fig. 2A, Fig. S2B). The surrounding somata were chosen randomly among the opsin-expressing cells, with a distance from the patched cell ranging from 58 µm to 268 µm (n = 110 surrounding cells) corresponding to an average distance of 141±8 µm (n = 11 patched cells; Fig. 2B). Similar patterns are used for connectivity mapping experiments. In particular, the patterns of ‘1 target’ and ‘10+1 targets’, both containing the patched cell, will be used to characterize the AP properties upon single-cell and multi-cell on-target stimulation, while the pattern of ‘10 targets’ will be used to provides indications of the spatial selectivity of the multi-spot stimulation.

**Figure 2:**
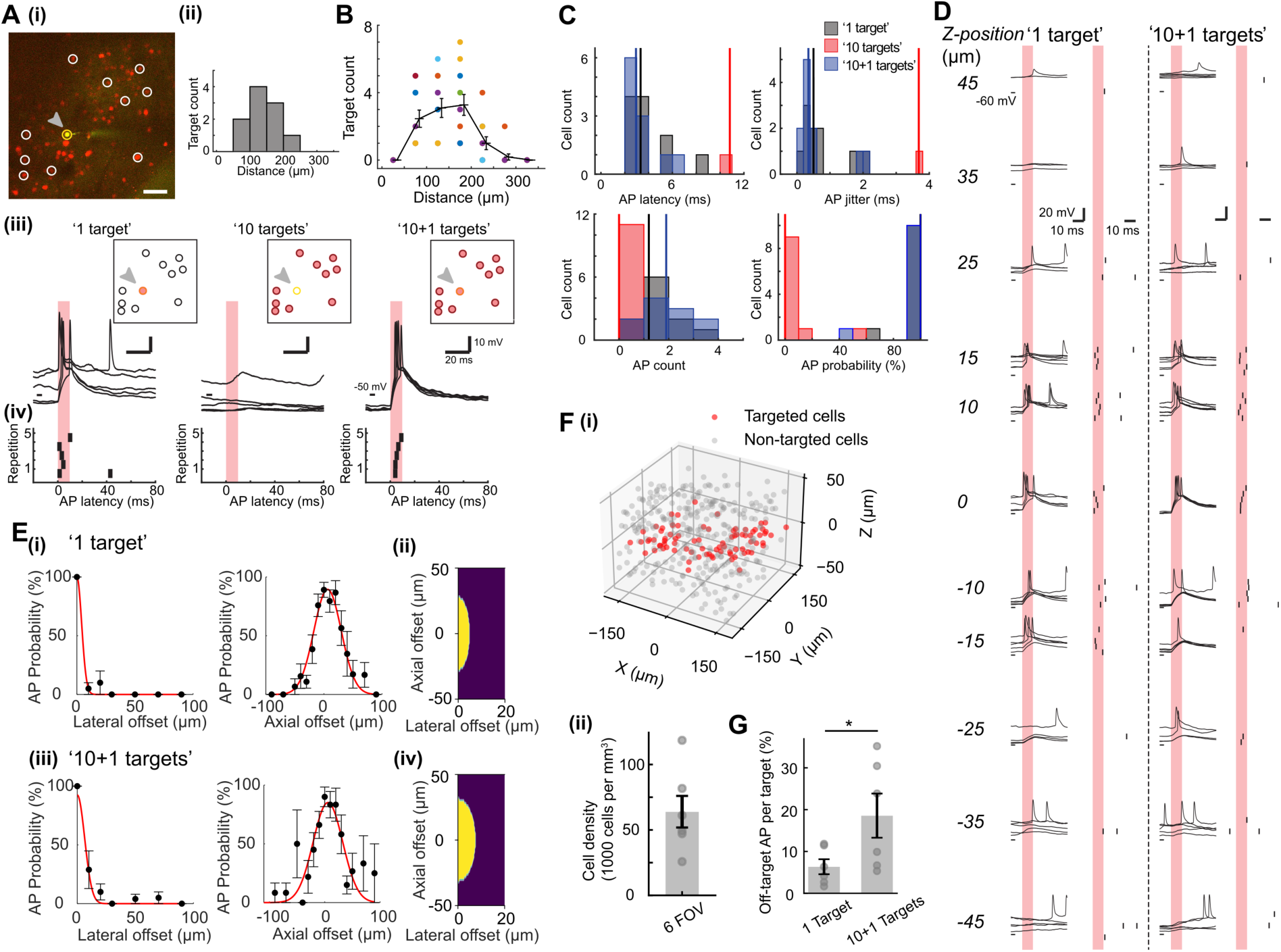
AP properties, spatial selectivity of AP generation, and estimation of off-target activation upon 2P holographic stimulation of single and multiple cells. (A) (i): Based on the fluorescence image from a defined FOV (scale bar 50 µm), one opsin-positive cell is selected for electrophysiological recording (yellow fluorescence from colocalization of opsin and Alexa-488 diffused from patch pipette, indicated by the yellow circle and the grey arrow) and 10 surrounding opsin-positive cells (red cells in white circles) are chosen to generate three holographic illumination patterns of ‘1 target’, ‘10 targets’, ‘10+1 targets’ (panel (iii)). (ii): Histogram of the distance between the 10 surrounding cells and the patched cell in this experiment. (iii): The three holographic patterns used to stimulate the corresponding targets (red shaded circles) while recording the membrane potential trace from the patched cell (grey arrow); excitation density 0.2 mW/μm^2^; 10 ms pulse, repeated 5 times. (iv): Raster plots of the corresponding AP peak latencies from the traces in panel (iii). (B) Spatial distribution of the spots in the ‘10 targets’ light-patterns. For each pattern, the distance between each of the target and the patched cell was distributed in one of the 50-µm bins between 0 and 350 µm. Colored dots indicated the bin counts in individual patterns. The black lines indicated the average bin counts and their s.e.m. for 11 patterns of ‘10 targets’. (C) Histograms of AP latency, AP jitter, AP count, and AP probability upon stimulation of ‘1 target’ (in black), ‘10 targets’ (in red), and ‘10+1 targets’ (in blue) (excitation density 0.15 - 0.3 mW/µm^2^; pulse duration 10 ms). Medians denoted as vertical lines of the corresponding colors (n=11 patched cells). (D) Examples of electrophysiological traces recorded in whole-cell configuration using 2P holographic stimulation with ‘1 target’ and ‘10+1 targets’ at different axial offsets relative to the patched cell. For each axial offset, 5 overlaid sweeps of traces with their raster plots of AP peak latencies are shown (excitation density 0.2mW/µm^2^; pulse duration 10 ms). Stimulation duration indicated as red shades. (E) AP probability as a function of the lateral and axial position of the ‘1 target’ (i) and ‘10+1 targets’ (iii) excitation patterns (n= 4 cells for lateral selectivity and n=10 cells for axial selectivity upon ‘1 target’ stimulation; n=5 cells for lateral selectivity and n=8 cells for axial selectivity upon ‘10+1 targets’ stimulation; mean±s.e.m.). Illumination condition: 0.2 mW/μm^2^,10 ms pulse duration. Red lines are gaussian fits. (ii,iv): Binary spiking probability function (SPF) upon ‘1 target’ and ‘10+1 targets’ stimulation. (F) (i) Example of the distribution of opsin-expressing cells within a volume of 350 µm x 350 µm x 100 µm, non-targeted cells (gray dots) an….d a set of coplanar targeted cells (red dots) at z≈0. (ii): Measured cell density across 6 FOVs (mean±s.e.m.). (G) Estimation of off-target AP generation probability per spot upon ‘1 target’ or ‘10+1 targets’ stimulation, by applying the SPF in (E) to the 6 FOVs reported in (F). Error bars mean±s.e.m. (p-value: 0.016, denoted by an asterisk, one-sided Wilcoxon signed-rank test)

By using single-cell illumination (‘1 target’), we investigated AP properties for power densities in the range of 0.1-0.6 mW/µm^2^ and illumination durations of 2-5 and 10 ms. A first characterization showed that photoactivated AP was achievable in 33/42 = 78.6% of the probed neurons when using stimulation power ≤0.3 mW/µm^2^. To maximize the AP probability in photostimulation experiments, we used illumination duration of 10 ms. Upon stimulation of increasing power density, we observed trends of increasing AP count and probability, and decreasing AP latency and jitter (Fig. S2B). We compared the AP properties upon 10 ms stimulation at different ranges of power density, here < 0.15 mW/µm^2^, 0.15-0.3 mW/µm^2^, > 0.3 mW/µm^2^ and found that power density significantly modulated AP probability (p=0.04) but not AP latency, jitter, count (p=0.59, 0.80, 0.28; one-way ANOVA for the 3 power density ranges, Table S1). For the connectivity experiments we chose a power density in the range of 0.15-0.3 mW/µm^2^ and a duration of 10 ms that across the full investigated samples induced an AP latency of 5.09 ± 0.38 ms (n=29 cells), AP jitter of 0.99 ± 0.14 ms (n=29 cells), AP count/pulse of 1.08 ± 0.11 (n=37 cells), and AP probability of 81.13 ±5.34% (n=37 cells). This choice of stimulation parameters therefore minimized the variability of presynaptic activation, contributing to the robustness of the connectivity mapping approach.

Next, using the same parameters defined above, we performed a second set of experiments to compare photostimulation properties under single- and multi-spot illumination. We observed induced APs upon stimulation patterns of ‘1 target’ and ‘10+1 targets’, but only rarely upon stimulation of ‘10 targets’: 1 AP in 45% of the case; 2, 3 AP in 34% and 14% of the case, respectively, for ‘1 target’ plus ‘10+1 targets’; 1 AP in 5% of the cases for ‘10 targets’ (a total of n=133-135 stimulations per type of pattern delivered over 11 cells inducing: 1, 2, 3 AP in 68, 47, 10 out of 133 stimulations for ‘1 target’; 1,2,3 AP in 52,45,27 out of 135 stimulations for ‘10+1 targets’; 1 AP in 7 out of 133 stimulations for ‘10 targets’) (Fig. 2C, Table 2), providing a first indication of the selectivity of the stimulation. Importantly, AP latency, jitter, AP count, and AP probability did not significantly vary upon single-cell and multi-cell stimulation (p=0.24, 0.38, 0.34, and 0.82 for AP latency, jitter, count, and probability, one-way ANOVA between n=11 cells under ‘1 target’ and ‘10+1 targets’ stimulation).

**Table 2:** AP properties upon 2P holographic stimulation of single and multiple cells of the working stimulation condition.

|  | AP latency (ms) | AP jitter (ms) | AP count | AP probability (%) |
| --- | --- | --- | --- | --- |
| '1 target' | 3.94±0.53<br>(n=11) | 0.83±0.21 (n=11) | 1.51±0.21 (n=11) | 95.45±3.66 (n=11) |
| '10 targets' | 10.82 (n=1) | 3.68 (n=1) | 0.061±0.047<br>(n=11) | 6.06±4.65 (n=11) |
| '10+1 targets' | 3.27±0.43<br>(n=11) | 0.60±0.21 (n=11) | 1.78±0.26 (n=11) | 93.64±5.44 (n=11) |

Data were presented as mean±s.e.m.

### Spatial selectivity of AP generation upon single-cell and multi-cell stimulation

Next, we quantified the spatial selectivity of AP generation upon single-cell and multi-cell stimulation by measuring the photoinduced spiking probability while shifting axially and laterally the 3 stimulation patterns (‘1 target’, ‘10 targets’, or ‘10+1 targets’) (see Methods) using the stimulation parameters in the range identified in the previous section (0.2 mW/µm^2^ power density and 10 ms pulse duration).

Upon ‘1 target’ or ‘10+1 targets’ stimulation, where the patched cell was targeted, we recorded AP probability of > 80% at zero lateral or axial offsets and progressively decreased when laterally or axially shifting the illumination pattern (Fig. 2D, E). The Gaussian fits of the spatial dependencies of the AP probability were used to derive the physiological resolution defined as the FWHMs of the curves (Lateral ‘1 target’: 9.6 µm, n=4 cells; Lateral ‘10+1 targets’: 14.9 µm, n=5 cells; Axial ‘1 target’: 55.5 µm, n=10 cells; Axial ‘10+1 targets’: 62.9 µm, n=8 cells) (Fig. 2E).

These values are consistent with the trends obtainable by convolving the (anisotropic) spot profiles (Fig. 1B(ii)) with the cell body profile and taking into account the dependence of photocurrent on excitation power (see Methods and Fig. S3). It is important to note that, in contrast to the case of the lateral “top hat” profile, the Gaussian profile along the axis produces tails outside the cell boundary, which significantly broadens the resolution at higher powers giving rise to the observed asymmetry of the physiological resolution.

Similarly to what has been done in ^69^, we used the values of the physiological resolution to estimate the 3D spiking probability function (SPF) defined as a binary function with lateral and axial size corresponding to the respective AP probability FWHMs for both single-cell and multi-cell stimulation (Fig. 2E(ii,iv)). Opsin-expressing neurons located within such ellipsoidal SPF were considered co-activated, while those outside were neglected. This choice was justified by considering that the lower light intensities on the periphery of the excitation volume would at most induce APs with lower reliability and of longer latency. As explained in the next paragraphs, this spiking activity is typically negligible within the time window during which the postsynaptic currents are analysed.

To estimate the corresponding density of opsin-expressing neurons, we used automatic cell recognition with Cellpose software ^81^ and manual adjustment afterwards to extract the coordinates of all opsin-expressing neurons within a volume of 350 µm x 350 µm x 100 µm of 6 representative FOVs. We found opsin-expressing cell densities ranging from 25800 to 118000 cells/mm^3^ (63.9 ± 12.1 *10^3^ cells/mm^3^, 6 FOVs) (Fig. 2F(i), gray dots). We then used this distribution to estimate the off-target AP probability of both single-cell and multi-cell stimulation, by counting the number of positive cells contained within the single-cell and multi-cell SPFs centered on each opsin-expressing neuron in the 6 FOVs at z ≈ 0 (Fig. 2F(i), red dots). We found that on average, the off-target AP probability (*P_OT_*) for single-cell stimulation was 6.4% ± 1.8% across the 6 considered FOVs (Fig. 2G), comparable to estimations from similar experiments in acute brain slices ^69,70^ (Table S3). For sparse connections (< 10%), this off-target activation would result in an overestimation of the connectivity rate values by approximately 0.6% and, even in the denser (*P_OT_* ≈12%, Fig. 2G), it would not exceed 1.2% (Fig. S4, see also similar estimations in ^69,70^. For multi-cell stimulation, the broader SPF gives rise to a larger *P_OT_* which increased to 18.6% ± 5.3% per spot (p=0.016, off-target AP generation upon ‘1 target’ vs ‘10+1 targets’ stimulation, n=6 FOVs, one-sided Wilcoxon signed-rank test) (Fig. 2G).

### In vivo connectivity mapping by using 2P holographic stimulation

Two strategies were employed for *in vivo* connectivity mapping: 1) *sequential connectivity mapping* where we sequentially activated single potential presynaptic cells (Fig. 3, S5A); 2) *parallel connectivity mapping* where we sequentially activated sub-groups of multiple potential presynaptic neurons (Fig. 4, S5A). In the latter case, the contributions of each presynaptic cell to the individual postsynaptic responses were reconstructed by using CS algorithms (Fig. S5B), as explained in the following paragraphs.

**Figure 3:**
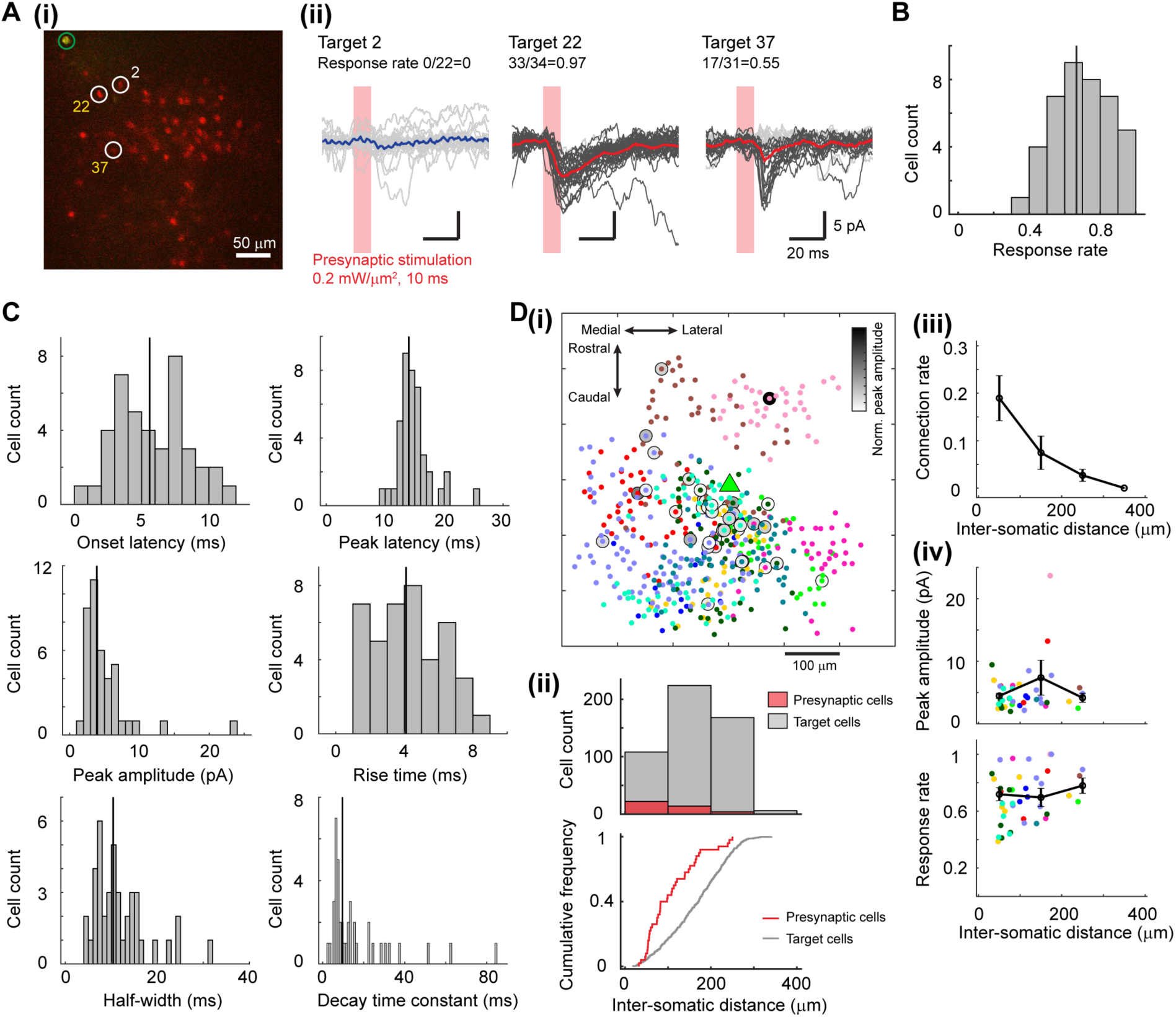
Sequential connectivity mapping by using single-cell stimulation. (A) (i): Based on the fluorescence image from a defined FOV (scale bar 50 µm), one opsin-negative cell is selected as the post-synaptic cell for electrophysiological recording (green cell indicated by the green circle) and each of the opsin-positive cells (red cells and examples indicated by white circles) is sequentially stimulated. (ii): Example traces of postsynaptic current in response to multiple stimulations of a non-connected cell (Target 2) and two connected cells (Target 22, Target 37). Non-responding individual traces denoted in light grey and the responding ones in dark grey. The corresponding average curves are reported in blue (Target 2) and red (Target 22 and 37). Excitation light density 0.2 mW/µm^2^; pulse duration 10 ms. The presynaptic stimulation periods are indicated by red shades. (B) Distribution of the response rates of the postsynaptic response upon single-cell stimulation of the presynaptic cell. Median denoted as vertical line (n=41 connected cells in 12 experiments). (C) Distributions of the properties of postsynaptic current in response to presynaptic stimulation of a connected cell. The response properties were extracted from the average traces across individual postsynaptic current traces. Medians denoted as vertical lines (41 connected cells in 12 experiments). (D) Spatial arrangement of probed neurons and identified connections. (i): Overlay of connection maps obtained in 11 experiments by using single-cell stimulation. The 11 postsynaptic cells were aligned at the center and denoted as a green triangle. Potential presynaptic cells stimulated in each experiment denoted as colored dots. Connected presynaptic cells indicated with grey shades, whose grey levels denoting the normalized peak amplitudes across all 40 connections. (ii): Histogram and cumulative frequency plot of inter-somatic distance between presynaptic and postsynaptic cells for targeted cells and connected cells (506 targeted cells, 40 connected cells, 11 experiments). Bin-size for histogram: 100 µm, n=108, 224, 168, 6 potential presynaptic cells and n=22, 14, 4, 0 connected cells in the 4 bins of 0-400 µm inter-somatic distance between targeted and postsynaptic cells. (iii): Connection rate as a function of inter-somatic distance between presynaptic and postsynaptic cells. For each experiment, the connection rate for each 100-µm bin from 0-400 µm was calculated as the number of connections divided by the number of stimulated cells within the same bin (mean±s.e.m., 11 experiments in total, n=11, 11, 11, 3 experiments in the 4 bins of 0-400 µm inter-somatic distance between connected and postsynaptic cells and between targeted and postsynaptic cells). (iv): Peak amplitude and response rate as a function of inter-somatic distance between presynaptic and postsynaptic cells. Peak amplitudes and response rates for connections in each experiment denoted as colored dots. The mean and s.e.m. of peak amplitudes and response rates of connections in each 100-µm bin indicated as black lines (11 experiments in total, n=8, 7, 4, 0 experiments in the 4 bins of 0-400 µm inter-somatic distance between connected and postsynaptic cells).

**Figure 4:**
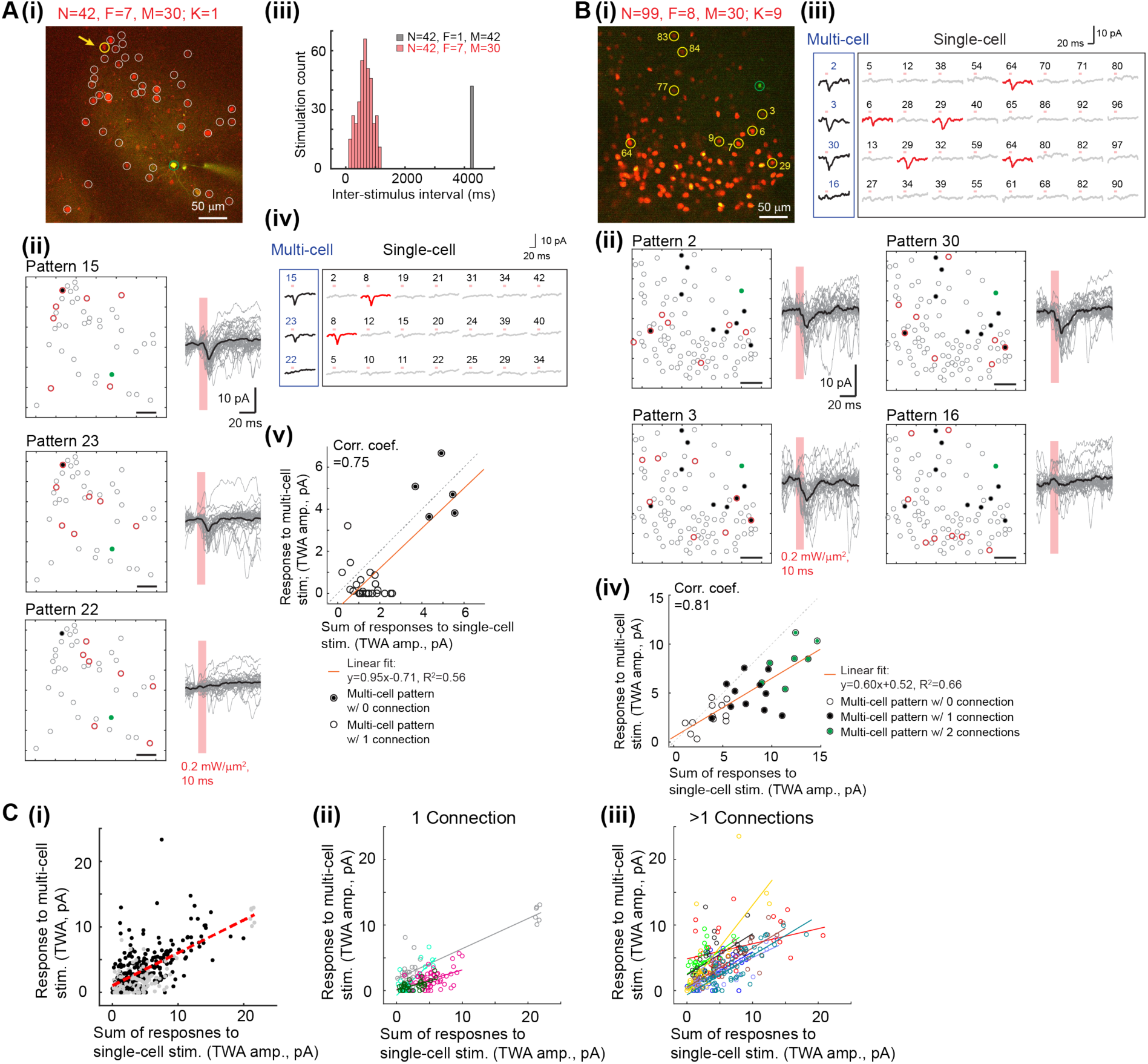
Postsynaptic activity in response to multi-cell stimulation. (A) Example of a multi-cell connectivity mapping experiment containing one presynaptic connected cell identified from the sequential connectivity mapping experiment. (i): Based on the fluorescence image from a defined FOV, one opsin-negative cell was selected for electrophysiological recording (green cell inside green circle) and N = 42 opsin-positive cells (red cells in white circles) were excited using M = 30 stimulation patterns each comprising F = 7 cells. The one connected cell (K=1) is indicated by a yellow circle and a yellow arrow. (ii): Example of three stimulation patterns, with the corresponding excitatory current traces. In the illustration of patterns, the postsynaptic cell was indicated in green, the potential presynaptic cells in grey open circles, the stimulated cells in red, and the 1 presynaptic cell (identified from the single-cell sequential mapping experiment) in black filled circle. The individual postsynaptic current traces are denoted in grey, and their average traces in black. Excitation density 0.2 mW/µm^2^; pulse duration 10 ms; 40, 35, and 32 repetitions for Pattern 15, 23, and 22 respectively. The stimulation pulses are in red shades, scale bar 50 µm. (iii): Histogram of the inter-stimulus intervals used in the stimulation protocols upon single-cell (F=1) and multi-cell (F=7) stimulation. (iv): Exemplary average current traces obtained using three stimulation patterns (of F=7 cells), together with traces corresponding to single-cell stimulation of the 7 cells targeted in these 3 patterns. The current trace upon single-cell stimulation of the connected cell denoted in red. (v): Scatter plot of the TWA amplitudes of the average current traces under multi-cell stimulation vs. the sum of those under single-cell stimulation. A linear fit (red line) and the correlation coefficient are reported. Patterns targeting 0, or 1 connected cell are indicated by empty, or black filled circles, respectively. (B) Example of a multi-cell connectivity mapping experiment using M=30 measurements each comprising F=8 cells from N=99 potential presynaptic cells, containing K=9 presynaptic connected cells (yellow circles) that were identified from the sequential connectivity mapping experiment. The post-synaptic cell is indicated by the green circle. Panels (i), (ii) the same as in (A)(i), (A)(ii); 29, 29, 30, and 28 repetitions for Pattern 2, 3, 30, and 16 respectively. Panels (iii), (iv) the same as in A(iv), A(v). In (iv), patterns targeting 0, 1 or 2 connected cells are indicated by empty, black filled or green filled circles, respectively. (C) Scatter plots of the TWA amplitudes of average current traces under multi-cell stimulation vs the sum of those under single-cell stimulation for: (i) all 12 experiments where 1 connection (data points in grey) or >1 connections (data points in black) were identified. (ii) 4 experiments where 1 connection was identified. (iii) 8 experiments where >1 connections were identified. In (ii) and (iii), colours encode for different experiments. Linear fits of either the full data set of all 12 experiments (i, red dashed line) or single experiments (ii and iii, coloured lines) are reported.

In both cases, in a FOV where L2/3 neurons expressed the somatic opsin ST-ChroME, we performed 2P-guided voltage-clamp recording to measure the excitatory synaptic current in a postsynaptic neuron (Fig. 3A, 4A, 4B). To prevent the generation of photocurrent from direct opsin activation, we recorded exclusively from opsin-negative neurons that were identified by the absence of red-fluorescence and a null photoinduced current upon soma illumination (Fig. S2C). Under the visual guidance of 2P imaging, we preferentially patched neuronal soma of pyramidal shapes, favoring postsynaptic recording from putative excitatory cells. Similar to what was discussed above, we verified that the patched neurons were likely putative excitatory cells according to their electrophysiological properties (opsin-negative cells in Table 1 and Fig. S2A). By holding the membrane potential at -70 mV, the reversal potential of inhibitory conductance, we recorded excitatory postsynaptic currents only, restricting our study to connections between putative excitatory neurons.

We extracted the coordinates of the N opsin-positive neurons surrounding the patched cell to define a set of potential presynaptic cells (Fig. S5A). We then designed accordingly a stimulation procedure consisting of M measurements, each corresponding to a stimulation with a distinct holographic light-pattern illuminating F targets. Under *sequential connectivity mapping*, *M = N* measurements of *F* = 1 target were sequentially used to probe each potential presynaptic cell one-by-one, whereas for *parallel mapping, typically M ≤ N* patterns were used to stimulate sub-groups of F > 1 cells (with F typically between 5 to 10, Table 3).

**Table 3:**
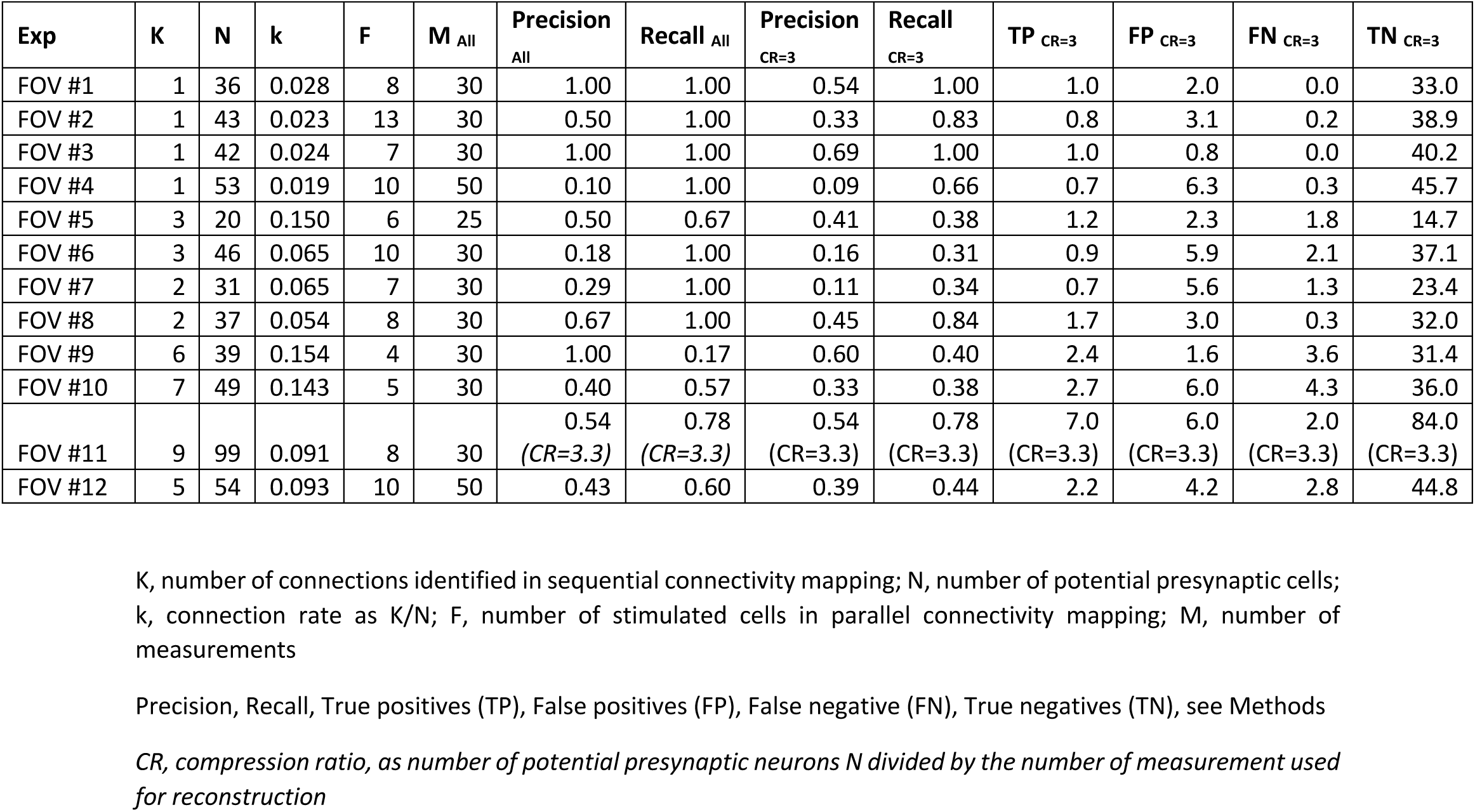
Results of sequential connectivity mapping and parallel connectivity mapping in 12 experiments.

| Exp | K | N | k | F | M<sub>All</sub> | Precision<sub>All</sub> | Recall<sub>All</sub> | Precision<sub>CR=3</sub> | Recall<sub>CR=3</sub> | TP<sub>CR=3</sub> | FP<sub>CR=3</sub> | FN<sub>CR=3</sub> | TN<sub>CR=3</sub> |
| --- | --- | --- | --- | --- | --- | --- | --- | --- | --- | --- | --- | --- | --- |
| FOV #1 | 1 | 36 | 0.028 | 8 | 30 | 1.00 | 1.00 | 0.54 | 1.00 | 1.0 | 2.0 | 0.0 | 33.0 |
| FOV #2 | 1 | 43 | 0.023 | 13 | 30 | 0.50 | 1.00 | 0.33 | 0.83 | 0.8 | 3.1 | 0.2 | 38.9 |
| FOV #3 | 1 | 42 | 0.024 | 7 | 30 | 1.00 | 1.00 | 0.69 | 1.00 | 1.0 | 0.8 | 0.0 | 40.2 |
| FOV #4 | 1 | 53 | 0.019 | 10 | 50 | 0.10 | 1.00 | 0.09 | 0.66 | 0.7 | 6.3 | 0.3 | 45.7 |
| FOV #5 | 3 | 20 | 0.150 | 6 | 25 | 0.50 | 0.67 | 0.41 | 0.38 | 1.2 | 2.3 | 1.8 | 14.7 |
| FOV #6 | 3 | 46 | 0.065 | 10 | 30 | 0.18 | 1.00 | 0.16 | 0.31 | 0.9 | 5.9 | 2.1 | 37.1 |
| FOV #7 | 2 | 31 | 0.065 | 7 | 30 | 0.29 | 1.00 | 0.11 | 0.34 | 0.7 | 5.6 | 1.3 | 23.4 |
| FOV #8 | 2 | 37 | 0.054 | 8 | 30 | 0.67 | 1.00 | 0.45 | 0.84 | 1.7 | 3.0 | 0.3 | 32.0 |
| FOV #9 | 6 | 39 | 0.154 | 4 | 30 | 1.00 | 0.17 | 0.60 | 0.40 | 2.4 | 1.6 | 3.6 | 31.4 |
| FOV #10 | 7 | 49 | 0.143 | 5 | 30 | 0.40 | 0.57 | 0.33 | 0.38 | 2.7 | 6.0 | 4.3 | 36.0 |
| FOV #11 | 9 | 99 | 0.091 | 8 | 30 | 0.54<br>(CR=3.3) | 0.78<br>(CR=3.3) | 0.54<br>(CR=3.3) | 0.78<br>(CR=3.3) | 7.0<br>(CR=3.3) | 6.0<br>(CR=3.3) | 2.0<br>(CR=3.3) | 84.0<br>(CR=3.3) |
| FOV #12 | 5 | 54 | 0.093 | 10 | 50 | 0.43 | 0.60 | 0.39 | 0.44 | 2.2 | 4.2 | 2.8 | 44.8 |

K, number of connections identified in sequential connectivity mapping; N, number of potential presynaptic cells; k, connection rate as K/N; F, number of stimulated cells in parallel connectivity mapping; M, number of measurements
Precision, Recall, True positives (TP), False positives (FP), False negative (FN), True negatives (TN), see Methods
CR, compression ratio, as number of potential presynaptic neurons N divided by the number of measurement used for reconstruction

### Sequential connectivity mapping

In all experiments, we used stimulation conditions (0.2 - 0.3 mW/µm^2^ intensity and 10 ms pulse duration) that, as discussed above, enabled AP generation with high reliability, millisecond latency and sub-millisecond jitter (Fig. 2C). We investigated 12 FOVs, corresponding to 12 postsynaptic neurons. Each FOV comprised 20-99 opsin-expressing cells resulting in an average number of N=45.8±5.6 probed presynaptic cells per experiment. The inter-somatic Euclidean distance between potential presynaptic and postsynaptic cells ranged between 18.2 and 341.0 µm (549 potential presynaptic cells in 12 FOVs). Various holographic phase masks, corresponding to the different illumination patterns, were precalculated and sequentially displayed on the SLM. The stimulation of each putative presynaptic cell was repeated 24-60 times in order to average postsynaptic currents across repetitions and assess their response rates. Specifically, in 9 experiments (FOV 1, 2, 5-11 in Table 3), holographic patterns were refreshed every 6 stimulation pulses delivered at 10 Hz, in other 3 experiments (FOV 3, 4, 12 in Table 3), they were refreshed at each stimulation pulse at a rate of 10 Hz and each cell was re-stimulated after sequential illumination of all other potential presynaptic cells, i.e., with inter-stimulus interval of N*0.1 s (see Methods). The latter protocol was preferable because the stimulation frequency could be kept below 1 Hz to avoid inducing potential short-term synaptic plasticity ^20,82^. Of note, individual postsynaptic traces with large fluctuations of spontaneous activity were discarded, resulting in 9-42 remaining repetitions per stimulation pattern (see Methods).

To eliminate from the measured postsynaptic currents the influence of polysynaptic contributions, spontaneous oscillations and the noise present in the whole-cell recording, the identification of post-synaptic current was based on two main criteria: 1) we set a peak amplitude threshold of 2-3 pA (corresponding to the typical recording noise level, Fig. S6B,C) on the averaged traces; 2) we performed a statistical test on the individual traces across repetitions, comparing current time-window averages (TWA, average current values in a specific time interval, Fig. S6B) few ms before and after photostimulation (see Methods for more details).

From 549 potential presynaptic neurons across 12 FOVs, we identified 41 connections (3.25±2.6 connections per FOV, ranging from 1 up to 9). Several representative examples of postsynaptic responses identified in planar or volumetric FOVs are presented in Fig. 3A and in Supplementary Fig. S6, S7, S8, S9. Overall, the identified connection rate, estimated as the ratio between responding sites in a FOV over total number of probed presynaptic cells, was of 7.6 ± 1.5%, ranging 1.9-15.4% across the FOVs. The sparse connection rate found here is in agreement with the connection rate values reported in previous *in vitro* studies for L2/3 excitatory neurons in mouse visual cortex ^15,16,68,70^.

From each identified connection, we then characterized the response rates (Fig. 3B), amplitudes, and kinetics (peak latency, rise and decay time constant, and current width; Fig. 3C, S8, S9, and Table S2). For the average current traces, the identified synaptic responses displayed peak amplitudes of few pA (5.0±0.6 pA, with the strongest detected response of 24 pA), fast rise time (4.3±0.3 ms) and short onset latency (5.8±0.4 ms). Interestingly, the latter, close to the photoinduced presynaptic AP latency (3.94±0.53 ms, Table 2), indicated a short synaptic delay and supported the monosynaptic nature of the identified connections ^14,20,83,84^. Additionally, in the following paragraphs, the magnitude of postsynaptic current responses will also be quantified from the current TWA between 10 and 15 ms after stimulation onset (typical time-interval for the occurrence of responses, Fig. S6B). The distributions of TWA amplitudes from connected and non-connected cells were found to be largely non-overlapping and values from the latter group approached zero (Fig. S6D), confirming the accuracy of our identification of connections and supporting the use of this TWA as a reliable proxy for quantifying responses.

The response rate of each connection, defined as the probability of detecting synaptic current across individual traces, was 0.70 ± 0.027 (ranging 0.39 - 1.00). This value, combined with the high probability in the presynaptic AP generation over multiple stimulation (previously characterized as > 90%, Fig. 2, Table 2) provides a good estimate of the synaptic failure rate. In agreement with previous reports, few connections of large amplitudes and high response rates were identified (as in Fig. S9) ^13,20,84^; such rare, strong and reliable connections may exert a dominant effect in network information processing ^9,85^. Occasional occurrence of double peaks in postsynaptic current responses (Fig. S9) may indicate either activation of 2 synapses or the occurrence of 2 presynaptic spikes of the target cell. Alternatively, it may be attributed to the unintended off-target activation of an additional presynaptic neuron together with the target one, as discussed above.

### Connection properties vs. distance between presynaptic and postsynaptic cell somata

Next, we analysed the dependence of the connection rate, strength, and the response with the distance between presynaptic and postsynaptic cell somata. Here we report results from presynaptic somata located in layer 2/3 and almost coplanar with the post-synaptic patched cell. Inter-somatic distance between postsynaptic and potential presynaptic cells was ranging from 18.2 µm to 341.0 µm (506 potential presynaptic cells over 11 FOVs). An example of detection of connection over multiple planes is reported in Fig. S7. Out of the 40 identified presynaptic cells, most of them (36/40 = 90%) were located within 200 µm from the patched neuron and no connections were identified above a distance of 250 µm (Fig. 3D(i)). To quantify this dependence, we analysed the connectivity vs. inter-somatic distance between presynaptic and postsynaptic cells by counting for each FOV the ratio of connected versus probed cells in circular crowns, R=100 µm thick, of internal radius r =n*R* (with n ranging from 0 to 4) centred at each postsynaptic cell in the FOV (Fig. 3D(ii)).

We found that the connection strength, estimated as the peak amplitudes of the average postsynaptic currents, did not co-vary with the inter-somatic distance (correlation coefficient=0.18, p=0.27, 11 FOV) (Fig. 3D(iv)). In addition, we observed that while the response rates were positively correlated with the inter-somatic distance (correlation coefficient=0.31, p=0.05, 11 FOV), connections of more variable response rates could be found at smaller inter-somatic distance.

### Parallel connectivity mapping by using multi-cell holographic stimulation and compressive sensing

To increase the throughput of connectivity mapping, we tested a *parallel* approach where we activated sub-groups of putative presynaptic neurons simultaneously and applied compressive sensing (CS) to reconstruct connectivity from the evoked postsynaptic response ^72,74^. CS gives conditions under which a system of equations can be solved with fewer measurements than unknowns. CS assumptions requires sparsity of connectivity, incoherently sampled measurements and linear summation of inputs. Based on the results from sequential connectivity mapping, the average connectivity rate of ∼8%, which decreased with inter-cell distances (Fig. 3D) indicated that our experimental conditions satisfy the sparsity condition for CS application. Incoherent sampling of measurements could be achieved by using sequences of multispot holographic patterns, ensuring that sufficiently distinct sub-groups of potential presynaptic neurons are stimulated in each trial ^76,86^. Regarding the linearity of input summations, one can expect both linear ^87,88^ and nonlinear summation ^89,90^. This dependence must therefore be verified in each experiment in order to enable the use of CS, as discussed below.

### Network reconstruction using compressive sensing: simulations, metrics and parameters

To design the protocol for patterned stimulation, we simulated a network of Izhikevich model neurons ^91^ with a user-defined connectivity rate *k* and a realistic distribution of synaptic weights and excitatory/inhibitory population, similar to ^74^. We used N potential presynaptic cells and mimicked the multi-cell optical stimulation protocol by selecting M different patterns of F excitatory cells that are simultaneously stimulated, i.e. we used F number of spots, on each trial. We used peak postsynaptic current responses as features in a basis pursuit CS algorithm to infer the estimated connectivity and compared them to the known connectivity in the model (see ^74,92^ and Methods).

Briefly, the CS algorithm aims to minimize over *x* the quantity 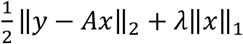, with *y* expressing the *Mx1* measurement vector (i.e. evoked postsynaptic current under patterned stimulation), A is the *MxN* measurement matrix (describing the stimulation patterns), ‖. ‖_1_ the *ℓ*_1_norm of the unknown *Nx1* connectivity vector *x* (i.e. synaptic strength) and λ is a regularization hyperparameter that balances sparsity of the solution and overfitting (Fig. S12). The 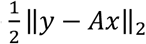 term corresponds to the error between the measured postsynaptic current and the linear sum of the weights connecting the stimulated cells to the postsynaptic cell on each trial. The λ‖*x*‖_1_ term is proportional to the absolute sum of the entries in *x* and acts as a penalty term that punishes dense, overfit solutions. To cover the experimental conditions obtained in the 12 FOVs investigated experimentally, simulations were performed over two different population sizes, (N=30, 100) and two connectivity rates (6.7%, 10%) corresponding to a total average number of connected cells, K, equal to 2 and 10, respectively.

Correct detection of connection presence or absence that matched the model’s ground-truth were declared true positives (TP) and true negative (TN), respectively. Similarly, incorrect detection of the presence of a connection or its absence were declared false positives (FP) and false negatives (FN) respectively (Fig. S10A). Both simulated and, later on, experimental reconstruction performances were quantified by monitoring 1) *precision*, defined as the ratio TP/(TP+FP), which indicated the fidelity of reconstruction; 2) *recall*, defined as the ratio TP/(TP+FN), which indicated the ability to detect the presence of all existing connections; and 3) *accuracy*, defined as the ratio ((TP+TN)/N), which specifies overall percentage of correct detection. In particular, we simulated reconstructions for different values of F (from 5 to 30% of N) corresponding to different coverages of the population (number of patterns that stimulate each cell, or *FxM/N*).

The results of the simulations, reported in Fig. S10B-E, provide a first glimpse of the feasibility of reconstruction, showing accuracy of up to 96% with a >2 fold reduction in the number of measurement compared to those used under sequential single-cell stimulation.

In accordance with previous predictions ^72,74^, we observed that, under low connectivity rate (k ≤10%), higher F values, and therefore better coverage of the population, improved reconstruction performances (Fig. S10B-E). Overall, given the limited improvement observed for F higher than 20% and in order to limit the degradation of the photostimulation spatial precision induced by high F values (Fig. 2E-G), we have designed the experimental protocol for parallel CS connectivity using stimulation patterns with an average number of cells F ranging between 4 to 13, corresponding on average to 19±7 % of the investigated presynaptic populations N. The chosen values for F also guaranteed that, even with 3-fold reduction in the number of measurements, each cell is typically targeted by at least 2 patterns, corresponding to a double coverage of the full population.

### Postsynaptic activity in response to patterned stimulation

In the same 12 FOVs discussed in Fig. 3B-3C and under analogous excitation conditions (power density 0.2 mW/µm^2^ and pulse duration 10 ms), we performed patterned stimulation (Fig. 4, S11) of 25 to 50 patterns for each FOV. According to the simulation described above, each of the pattern targeted on average F= 8±2.5 spots per pattern (ranging from 4 to 13), out of the N potential presynaptic cells, ranging between 20 and 99 cells. Each multi-cell stimulation pattern was repeated 18-60 times, refreshing the SLM pattern either every single pulse or every series of 6 pulses, as describe for sequential stimulation (see details in Methods). Even in this case, the former protocol is favorable as it allows to decrease stimulation frequency of the same target and, despite the more recurring illumination of cells under multispot excitation, this frequency could be kept below few Hz (Fig. 4A(iii)), beneath the threshold for inducing plasticity effects (Jouhanneau et al., 2015; Ko et al., 2013). After discarding individual traces with large fluctuation of spontaneous current activity, 6 to 43 repetitions, i.e., 31%-100% of the total repetitions, were further analysed.

Upon stimulation of multiple potential presynaptic cells, the postsynaptic cell displayed excitatory current of short onset latency, (Fig. 4A(ii), 4B(ii), Fig. S11), resembling the synaptic current responses upon presynaptic spiking (Fig. 3A(ii)). Criteria for inclusion of postsynaptic currents upon stimulation of 210 out of 395 multi-cell patterns were identical to those under single-cell stimulation (see Methods). The higher probability of observing postsynaptic responses, at 53% in the case of multi-cell stimulation, compared to the previously obtained 8% connection rate, hinted multi-synaptic activations and synaptic summations.

To quantitatively examine how the postsynaptic neuron integrated multiple presynaptic inputs, we calculated the TWA amplitudes from the averaged postsynaptic current traces for both single-cell and multi-cell stimulation (see Methods). Since we focused on the integration of excitatory synaptic inputs, negative TWA amplitudes (equivalent to outward currents), possibly from recording noise, spontaneous activity, or remaining inhibitory inputs, were offset to zero. For each multi-cell pattern stimulation, its TWA amplitude was compared with the sum of those calculated from the averaged response traces obtained in single-cell stimulation trials (Fig. 4A(iv,v), 4B(iii,iv)).

For the 12 FOVs investigated, the TWA amplitudes corresponding to multi-cell stimulation were positively correlated with the sum of those corresponding to single-cell stimulation (average correlation coefficient, on the 12 FOVs, = 0.66±0.05, ranging 0.37-0.91; p=0.045, 0.0064, 0.0012 for 3 experiments and p<0.001 for 9 experiments). We extended the same analysis by splitting the 12 FOVs into 2 groups, those containing 1 connected cell, where no summation was expected under multi-cell stimulations (K=1, 4 FOVs), and those containing > 1 connected cells (K>1, 8 FOVs). In all FOVs, we measured a positive correlation (average correlation coefficient, on the 12 FOVs, 0.68±0.10, ranging 0.45-0.91; p=0.0012 for 1 experiment and p<0.001 for 3 experiments in the first group; coefficient=0.65±0.06, ranging 0.37-0.81; p=0.045, 0.0064 for 2 experiments and p<0.001 for 6 experiments in the second group).

The slopes of the linear fits of the TWA upon multi-cell stimulation vs. the sum of those upon single-cell stimulation were of 0.59±0.09, ranging from 0.23 to 1.28 (n=12 experiments). By merging all the 12 FOVs together, we obtained a linear fit with a slope of 0.49 (395 multi-cell patterns in the 12 experiments) (Fig. 4C(i)), while the average slopes across experiments were 0.50±0.16, ranging 0.28-0.95 in the pool of experiments with 1 connected cell (Fig. 4C(ii)), and of 0.64±0.11, ranging 0.23-1.28 in the case of > 1 connected cells (Fig. 4C(iii)). In general, we noticed a large variability across FOVs for the correlation coefficients and slopes of the linear fits. In particular, several FOVs showed linear fits with slopes <1 indicating that the TWA amplitudes under multi-cell stimulation were smaller compared to the sum of those under single-cell stimulation (12 experiments). Such sub-linearity might have different explanations. It could result from increased access resistance during measurements of multi-cell stimulation (in 8 from the 12 experiments, measurements of multi-cell stimulation were performed *after* those of single-cell stimulation), which decreases current amplitudes. Additionally, it could be attributed to the sublinear dendritic integration of closely located synaptic inputs, where a localized depolarization within the dendritic branch may result in a reduction of the driving force, as previously observed in cortical pyramidal neurons ^93–96^. Finally, a potential role could be played by the photostimulation of inhibitory neurons that, given the known dense local inhibitory connectivity ^29,70,97,98^, could decrease the photoactivation of target cells and, consequently, reduce the detected responses on the patched cell ^46,99^.

Nevertheless, besides the absence of a perfect summation, the overall positive correlation between multi-cell responses and the summation of single-cell responses supported the application of compressed sensing strategies.

### Reconstructing connectivity by using compressive sensing

We then tested whether responses recorded under multi-cell stimulation can be demixed to identify connected cells using an adapted basis pursuit CS algorithm (see Methods). In the 12 FOVs investigated (Fig. 3B-3C, Fig. 4C), responses were reduced to scalar values considering the TWA amplitudes of averaged postsynaptic current traces as inputs to the reconstruction algorithm. K-mean clustering for 2 clusters were performed to establish a connection-defining threshold to determine whether the reconstructed responses of all potential N presynaptic cells were classified as connected or non-connected cells. (Fig. 5A, 5B, Table 3). The performance of the reconstruction was evaluated according to precision and recall (Fig. S10A). For each FOV, we assigned a certain number of multi-cell stimulation patterns M, varying between 25-50, corresponding to approximately 0.3 -1.25*N, determined according to the estimated experiment duration, mainly limited by the possibility to maintain good whole-cell recording over the full sequence. We then performed CS reconstruction using all or some of the measurements available, exploring reconstruction performance using different values of compression ratio (CR, defined as N/M).

**Figure 5:**
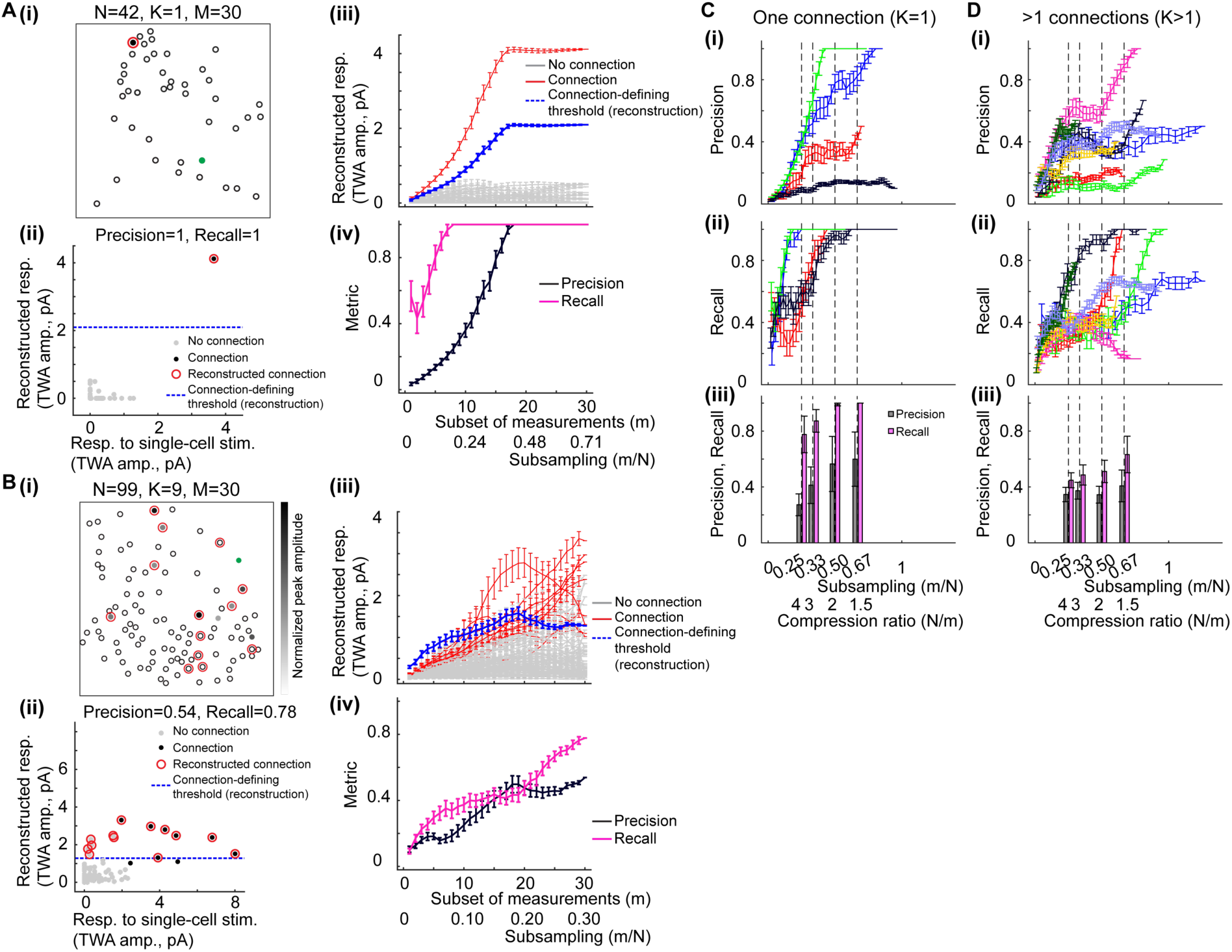
Reconstructing network connectivity by using compressive sensing. (A) Example of parallel connectivity mapping in a FOV where 1 connection was identified by using sequential connectivity mapping, the same example as in Figure 4A. (i): Map of the coordinates indicating the postsynaptic cell (green filled circle), potential presynaptic cells (black open circles), identified presynaptic cell (filled circle in black), and the reconstructed connection by using all measurements of multi-cell stimulation (red open circle). (ii): Scatter plot of the reconstructed responses using compressive sensing vs. the ground-truth TWA responses (TWA amplitudes) from single-cell stimulation traces, for the N=42 potential presynaptic cells. To identify connections after reconstruction, a connection-defining threshold determined by k-mean clustering (see Methods) is denoted as blue dashed line. Black and grey dots identify connection and no connections, respectively, identified in sequential mapping; red open circles identify reconstructed connections. Precision and recall of the reconstruction results by using all 30 measurements were specified as plot title. (iii): Reconstructed responses for the 42 potential presynaptic cells and the connection-defining threshold (blue) as a function of the measurement number m, i.e. number of subsampled multi-cell stimulation pattern used for the reconstruction of the total M available. Subsampling indicate the ratio m/N. Identified connections in sequential connectivity mapping are indicated in red and identified no connections in grey. For each subsampling value (with m<M), the synaptic responses were reconstructed and averaged over a total of M random subsets of multicell to ensure balanced sampling across all potential presynaptic cells (see Methods). (iv): Precision and recall of the reconstruction result as a function of the measurement number in subsampling. Error bars as mean±s.e.m. (B) Example of parallel connectivity mapping in a FOV where 9 connections were identified by using sequential connectivity mapping, the same example as in Figure 4B. Panels (i-iv) the same as in panels A(i-iv). (C) (i,ii): Precision and recall as a function of the subsampling m/N for 4 experiments where 1 connection was identified in sequential connectivity mapping. Similar to A(iii-iv), precision and recall for each subsampling were assessed and averaged over M different subsets of multicell stimulations. Precision and recall of different experiments indicated as different colors. Error bars as mean±s.e.m. (iii): Summaries of precision and recall at different subsampling which corresponded to compression ratios of 4, 3, 2, 1.5 (n=4 experiments for each condition). Error bars as mean±s.e.m. (D) (i,ii): Same as C(i,ii) for 8 experiments where >1 connections were identified in sequential connectivity mapping. Precision and recall of different experiments indicated as different colors. (iii): Summaries of precision and recall at different measurement numbers which corresponded to compression ratios of 4, 3, 2, 1.5 (n=8, 8, 7, 6 experiments). Error bars as mean±s.e.m.

Using all the M measurements available per FOV, we obtained the following reconstruction performances: for the 4 experiments where sequential mapping identified one connection, we achieved a precision = 0.65±0.22, ranging from 0.1 to 1, and recall = 1.00±0 in 100% of the cases (with CR=1.27±0.09, ranging from 1.06 to 1.43); for 8 experiments where sequential mapping identified >1 connections, we obtained a precision = 0.50±0.09, ranging from 0.18 to 1, and recall = 0.72±0.10, ranging from 0.17 to 1 with CR=1.49±0.28, ranging from 0.8 to 3.3 (see Table 3).

To examine how measurement number affected the reconstruction performance, we computed the evolution of precision and recall as a function of the number of measurements (see Methods). We performed the reconstruction by selecting random subsets of *m* measurements out of the M measurements available. A decreased *m* corresponded to a lower subsampling (*m/N*), or a higher CR (N/m). Trends of increasing precision and recall were observed when the measurement number was increased, or when CR was decreased (Fig. 5C-5D). We focused our attention to the performance achievable under a compression ratio of CR=3, which resulted in reconstruction performances of precision=0.41±0.13 and 0.37±0.06 and recall=0.87±0.08 and 0.49±0.07 for experiment with 1 and >1 connections, respectively (Table 3), meaning that, besides the presence of several FP, always more than half of the connected cells were retrieved while, for the highest sparsity conditions of 1 connection, the single connected cell was identified in 87% percent of the cases.

The reported results were obtained with a fixed regularization factor λ=0.1. We also investigated the effect of lower or higher λ (Fig. S12A-D), preferring either recall (Fig. S12D) or precision (Fig. S12C) respectively. Similarly, the connection-defining threshold to identify positive cells could represent an additional adjustable parameter, such that decreasing its value could allow to decrease the number of missed connections (FNs) and increase TPs so to increase average recall values from ≈50% to 70% for FOVs with >1 connections at the price of higher number of FPs (precision from ≈40 to 20%) (Fig. S12 E). Overall, we note how better performances were achieved for experiments with single connections, in which both recall and precision could reach 1 (Fig. 5A, 5C, Table 3), meaning that the connection was retrieved, without FNs and FPs. As for experiments with >1 connections, FNs were still limited (1.4±1.3, mean±SD, always ≤3), while FPs were more frequent (4.7±4.3, mean±SD) (Fig. 5B, 5D, Table 3). This general behavior was in accordance with theoretical framework of compressive sensing and previous modeling studies ^72,74,100^, in which the required number of measurements was expected to scale linearly with connection number according to M α K log(N).

## Discussion

We have presented an approach for high-throughput *in vivo* connectivity mapping combining two-photon optogenetics with spatiotemporally shaped parallel stimulation and compressed sensing. We achieved optical control of presynaptic activation at cellular resolution by using 2P excitation, with temporally-focused holographic spots, of neuronal somata expressing the fast efficient soma-restricted opsin ST-ChroME ^66^ while the induced postsynaptic responses were detected using whole-cell recording. We demonstrated two methods for high-throughput connectivity mapping. In the first method, the connections between a chosen postsynaptic cell and *N* potential presynaptic cells, contained within a 350 µm x350 µm x100 µm volume, were probed through single-cell sequential activation of the *N* cells. In the second one, connections were probed in the same population of cells by replacing single-cell stimulation with parallel stimulation of subsets of potential presynaptic cells, each containing *F* cells ^72^. In the second case, the contributions of each presynaptic cell to the individual postsynaptic responses were reconstructed by using a compressive sensing algorithm.

The combination of temporally-focused holographic light-shaping and soma-targeted opsins allows the spatio-temporal precision that is essential for *in vivo* connectivity mapping. Specifically, confirming previous finding ^54,55,58,60,61,66^, we have shown that soma illumination enables the generation of reliable APs with few millisecond latencies and sub-millisecond temporal jittering and that the temporal resolution and precision were maintained when using multi-target illumination. Since mono-synaptic responses typically occur within < 5 ms after the presynaptic AP firing ^16,18,20^, precise knowledge of the presynaptic spiking time is essential to discriminate true synaptic responses from spontaneous or polysynaptic events.

Combining temporally-focused holographic excitation with the soma-targeted opsin ST-ChroME, we have demonstrated a lateral and axial spiking probability resolution of 10 vs 15 µm, and 55.5 vs 62.9 µm, for single-cell vs multi-cell stimulation, respectively. These values are the result of a trade-off between the use of sufficiently high power to ensure a high probability of spiking rate without compromising the spatial resolution. Future improvements in targeting strategies (e.g., through the production of transgenic lines with higher and more homogenous opsin expression) will make it possible to work at lower powers and thus improve the physiological spiking probability resolution. To further reduce the uncertainty due to possible off-target excitation, one could adopt previously proposed strategies based on volumetric calcium imaging around the target cell ^71^ or sequential activation of axially-shifted stimulation patterns ^68^, although this will add complexity to the experimental pipeline and reduce the accessible number of probed pairs.

Here, we have optimized an experimental pipeline including a fast GPU calculation of the photostimulation hologram, a fine-tuning of the number of repetitions (typically ≥ 30) and of the stimulation sequences (changing target at every photostimulation pulse), to improve the signal-to-noise ratio in the recorded responses and avoid potential opsin inactivation and plasticity effect. This allowed to probe 100 presynaptic sites in ∼5 minutes. A larger number of pairs could be probed by shortening even further the time interval, Δt, between two sequential stimulations. This is the sum of the SLM refresh time, t_SLM_, added to the longer time between the photostimulation time, t_ill_, and the electrophysiological trace recording time, t_rec_. That is Δt = t_SLM_ + t_ill_ or Δt = t_SLM_ + t_rec_ if t_rec_ > t_ill_. In our experimental conditions we used Δt ≈100 ms (with t_SLM_≈40 ms). The use of faster SLM (refresh rate 300Hz) ^42^, that will lower t_SLM_ down to ≈ 2-3ms, or the use of approaches for fast (50-90 μs) scanning through multiple holograms ^101^, also combined with current deconvolution algorithms ^75^, could reduce the sequential time interval to Δt ≈ t_ill_ or Δt ≈ t_rec_ if t_rec_ > t_ill_.

Single-cell sequential activation enabled to retrieve ∼ 8% of connected cells out of > 500 presynaptic neurons probed. This value needs to be compared with the values of 2-19% reported in the literature for *in vitro* studies of excitatory connections in L2/3 of mouse visual cortex, obtained either through multi-electrode recording or optogenetic approaches ^15,16,68,70^. Higher connectivity rate could be expected *in vivo* due to the absence of connection losses induced by slicing procedure ^4–6^. On the other side, contrary to previous study, here we might have induced opsin expression in both excitatory and inhibitory neurons, while voltage clamp at -70 mV only reveals excitatory inputs, which might therefore lower the estimated number of detected connections. Underestimation of connection number could also be a consequence of the non-uniform opsin expression in potential presynaptic neurons, so that a fraction of the target presynaptic cells (measured at ≈20% as ratio of the opsin-positive cells spiking under photostimulation), might not have responded reliably to photostimulation. Also, the noisier whole-cell recording *in vivo*, and the consequent strict condition we set to identify reliable synaptic response, could have precluded detecting weak connections (connection strength < 2-3 pA or with high synaptic failure rate).

Using multi-cell patterned excitation combined with compressive sensing algorithms, we have demonstrated that, depending on the compression ratio CR imposed on the measurement number, we could retrieve between 28 (CR=1.4±0.2) to 22 (CR=3) out of the 41 total connections identified by sequential mapping. The accuracy of reconstruction was particularly high in FOVs with sparse connectivity (≈2%) where a CR=3 was enough to retrieve the single connected cell with almost no missed connection of FN (recall >0.8) and at most 2.4 average FP per FOV over typically ≈ 45 cells (precision > 0.5). For FOVs with higher connectivity rate (≈10%), the approach missed an average of 2 connected cells per FOV and gave an average of false positive FP ≈ 4 (reaching up to FP = 6 in some cases). To ensure nearly unitary precision and eliminate FPs even in these extreme cases, the small set of retrieved connections can be verified through single-spot sequential stimulation, which necessitates only a minimal number of additional measurements (typically around 10-15% of N) and proportionally minimal reduction of the CR. Despite the inevitable missed connections, this approach will enable fast identification of subsets of presynaptic cells, particularly useful e.g. for investigation of synaptic plasticity or synaptic integration.

Fine-tuning the regularization parameter λ and the threshold values to identify positive cells could allow to further guide the reconstruction toward denser or sparser solution, reducing either FPs or FNs according to specific experimental needs.

Compressive sensing connectivity mapping will become particularly advantageous when exploring sparse connections (less than 10 ^14,19,20,102,103^) in populations comprising hundreds or thousands of potential presynaptic cells, a scale that is either inaccessible or highly impractical with single-cell sequential mapping. In such scenarios, efficient mapping requires striking a balance between the number and distribution of F spots per pattern which ensures comprehensive coverage of the entire volume (larger F) while minimizing off-target stimulation (sparse distribution of F targets).

So far, the stimulations parameters (F, M) and the distribution of the F targets within each of the M patterns used in each sequence were predefined before the experiment based on the expected connectivity rate.

The number of measurements M could be further decreased by implementing an active feedback control mechanism which enables the removal of the F_i_ cells belonging to the matrix row M_i_ (the *i-th* pattern) that do not elicit any current during stimulation from subsequent stimulation patterns.

Previous theoretical works ^72^ suggested that a main source of errors in the reconstruction is the missing or uncontrolled photostimulation events. Similar to the sequential mapping, the ability to monitor the activity of targeted cells and other neighbouring cells, i.e. combining opsins with an activity indicator, could allow to assess the responses of the target presynaptic cells ^69^, detect potential off-target stimulation ^71^ or monitor *in vivo* spontaneous activity in order to adjust the stimulation matrix M accordingly to ensure presynaptic cell states are stationary across different times.

Here, spiking probability and time invariance of presynaptic spike induction were characterized using patch-clamp electrophysiology. This step can be avoided by integrating in the experimental protocol with optical readout of presynaptic activity using GCaMP imaging ^58,69,71^ or using voltage indicators. In the latter case, the evoked presynaptic response as well as precise spike times can be verified ^104^ to correct for any jitter in synaptic inputs, which is key for accurate reconstruction ^74^.

In summary, this study successfully demonstrated *in vivo* optogenetics-based mapping of synaptic connectivity in the mammalian brain, achieving results that were previously unattainable. Notably, the capability to rapidly probe numerous connected cells per minute of recording facilitated the reconstruction of a detailed local map of excitatory connections within the L2/3 of the mouse visual cortex. This parallel mapping approach holds immense potential in further improving the efficiency of connectivity mapping in behaving animals, particularly for revealing long-range connections across spatially distinct brain regions characterized by sparse connectivity ^18,35,69,84,105–107^.

## Acknowledgments

We acknowledge support from the IHU FOReSIGHT (Grant P-ALLOP3-IHU-000), the Fondation Bettencourt Schueller (Prix Coups d’élan pour la recherche française), the Axa research funding (Axa Chair), the Région Île-de-France (Grant WASCO; DIM-Elicit), the France-Israel Scientific Research Program (”Maimonide-Israel” program), the Agence National de la Recherche and the European Research Council (HOLOVIS; ERC2019-ADG-885090 to V.E.) for financial support. The project has also received funding from the European Union’s Horizon 2020 research and innovation program under the Marie Skłodowska-Curie grant agreement #747598 (I.C.) and #813457 (C.Y.C.).

We would like to thank Kris Blanchard for preliminary test and implementation of mapping protocol in *in vitro* preparation, Valeria Zampini for help with viral injection and for useful discussions, Eirini Papagiakoumou for the help in the setting up of the optical system, Hillel Adesnik Lab for providing ST-ChroME opsin virus, Christophe Tourain for the support in the implementation and maintenance of the experimental apparatus, and Manuel Simonutti, Julie Degardin, and Quenol Cesar from the animal facility of the Institut de la Vision for their help and support with the animal experimentation.

## Data availability

All data are available upon request.

## Code availability

All codes used to process and analyse data are available upon request.

## Materials and methods

### Animals

All animal experiments were performed in accordance with Directive 2010/63/EU of the European Parliament and of the Council of 22 September 2010. The protocols (APAFIS#14267–201803261541580 v3) were approved by the author’s institutional ethics committee for animal research (CEED5). Experiments were performed by using adult male C57BL/6J mice (Janvier Labs). Some experiments of AP properties upon single-cell photo-stimulation were performed on adult female or male mice of transgenic line GP4.3 (The Jackson Laboratory) which expressed the calcium indicators GCaMP6s ^108^. Viral injection for expressing opsins in cortical neurons were performed on 4-week-old mice. Holographic stimulation experiments of measuring induced spiking activity or mapping synaptic connectivity were carried out on mice 3-12 weeks after injection.

### Viral injection and surgical procedures

Stereotaxic injection of viral vectors was performed when the mice were anesthetized with intraperitoneal injection of a ketamine-xylazine mixture (0.1 mg of ketamine plus 0.01 mg of xylazine per g body weight). Acute experiments involving holographic stimulation were carried out when mice were anesthetized with isoflurane (2% for induction and 0.5-1% for experiment). Viral vectors of AAV9-hSyn-DIO-ChroME-Flag-ST-P2A-H2B-mRuby3-WPRE-SV40 in combination with AAV9-hSyn-Cre-WPRE-hGH were used for obtaining soma-restricted expression of the opsins ST-ChroME. Through a craniotomy over V1 (3.5 mm caudal from bregma, 2.5 mm lateral from midline), 0.8-1 µL of viral vectors were infused via a cannula at 250 µm deep (L2/3 of V1) at a speed of 80-100 nL/min. To perform acute photo-stimulation experiments *in vivo*, the mouse was head-fixed by attaching the skull to a small metal plate. A circular craniotomy of 2 mm diameter was made over V1 and the dura mater was removed. Agarose of 0.5-2% and a cover glass were applied on top of the craniotomy to dampen the tissue movement.

### Optical system for 2P holographic stimulation

The custom-built optical system included two paths enabling to perform 2P holographic stimulation and 2P scanning imaging.

The configuration for generating 2P holographic light-patterns by using computer-generated holography with temporal focusing was described in ^48^. The light source of stimulation was provided by a fiber amplifier laser of 1030 nm and 250 fs pulse-width operating at a repetition rate of 500 kHz with maximum average power of either 10 or 50 W (Satsuma HP or HP3, Amplitude Systems). The stimulation laser first passed through a polarizing beam splitter to ensure that the polarization was linear, then the wavefront of the laser beam was modulated by passing through a static phase mask (Double Helix Optics) to generate a circular holographic light-spot. The phase mask profile was calculated according to Gerchberg-Saxton (GS) algorithm. The laser beam then passed through a Fourier lens (L5, f=190mm) and projected (L6-7, f=400mm and 400mm) on a blazed grating (830 lines/mm, Blaze Wavelength 800 nm) to generate the effect of temporal focusing. The diffracted first-order beam after the grating was collimated via a lens (L8, f=750mm) and then projected onto the sensitive area of a reconfigurable liquid crystal SLM (Hamamatsu Photonics). The SLM was controlled by Matlab (MathWorks) and a custom-designed software Wavefront IV ^109^, which implemented GPU-accelerated GS algorithm (adapted from ^110^) to generate one or more diffraction-limited targets to multiplex the holographic spot generated by the static phase mask. The size of the laser beam was adjusted by a telescope (L9,10, f=500, 750mm) to fill the back focal plane of the microscope objective (Nikon W APO NIR of 40X, or Nikon CF175 LWD of 16X). A hand-made zero-order block was placed at the intermediate image space after L9 to physically suppress the zero-order diffraction of SLM in the light-pattern. The triggering and power of stimulation laser were controlled by the internal control of the laser head in combination with custom-written Matlab script, a data acquisition system (National Instruments), a digitizer (Digidata 1550A, Molecular Devices), and Clampex software (Molecular Devices).

The configuration of 2P scanning imaging was similar to that described in ^61^. The light source of imaging was provided by a Ti:Sapphire laser (Coherent, Chameleon Vision 2), whose power was controlled by a liquid crystal variable phase retarder (Meadowlark Optics, LRC-200-IR1) and a polarizer tube (Meadowlark Optics, BB-050-IR1). The imaging was performed by raster scanning with a pair of galvanometric mirrors (3-mm aperture; Cambridge Technology, 6215H series). There are two telescopes before (L1,2, f=75,175mm) and after (L3,4, f=50,350mm) the galvanometric mirrors to fill the galvanometric mirrors and back focal plane of the microscope objective, respectively. The imaging and stimulation lasers were combined before the back focal plane of the microscope objective by using a dichroic mirror.

The emitted fluorescence signal from the sample was collected by the same microscope objective and reflected by a dichroic mirror. Direct reflection of laser was filtered by an infrared light-blocking filter (Semrock, FF01-750sp). The fluorescence signal was split into two channels (red and green) by a dichroic mirror (Semrock, FF555-Di-3) and two emission filters (Semrock, FF02-617/73 and FF01-510/84) and was collected by two photomultiplier tubes (Hamamatsu Photonics, R3896 and R9110 SEL). Imaging acquisition was controlled by using the software ScanImage (Vidrio Technologies).

### Calibration of the optical system

Spatial calibration between the stimulation and imaging fields was done to compensate for the small but non-negligible physical misalignment between the two optical paths, so that the stimulation light-spot(s) can precisely target the opsin-expressing neurons identified by using 2P imaging. To do that, first, we photo-bleached on a thin rhodamine layer with holographic light-spots in 2D and acquired a 2P image of the photo-bleached rhodamine layer. Next, the misalignment was estimated by applying stretching, rotation and translation in the stimulation field to match the targeted coordinates, i.e., the actual bleached positions. Axial displacement between imaging and stimulation focal plane were compensated by adding a phase-wrapped spherical lens function to the SLM.

To measure the inhomogeneity in the intensity of light-spot due to the diffraction efficiency loss of the SLM, one or more (5,10 and 15) holographic spots, within a 6×6 grid coordinates with small random offsets in a FOV of 350 µm, were sent to a 1-mm thick fluorescence slide (FSK6, Thorlabs) and the fluorescence images were collected by the same objective lens and imaged by a CCD camera (Hamamatsu C8484-05G02). In case of one holographic spot per pattern, the diffraction efficiency of each of the coordinates were sampled in one light-pattern, while in case of multiple holographic spots per pattern, each of the coordinates were sampled in 3-5 different light-patterns.

To compensate for the diffraction efficiency loss, we adapted the method described in ^55,111^. Briefly, the total power was adjusted by the inverse of total diffraction efficiency, and the weights of different light-spots within the same pattern, which determined the relative power ratio, were adjusted individually by the inverse of the diffraction efficiency of the respective spot. The diffraction efficiency profile of one spot per pattern was slightly different from that of multiple spots per pattern, and thus their compensations were carried out separately.

For the characterization of optical axial profile and visualization of 3D spots distributions, a thin rhodamine poly(methyl methacrylate) (PMMA) layer spin-coated on a glass slide (less than 1 µm in thickness) was imaged from below by using a sub-stage objective and a CMOS camera (MQ013MG-ON, Ximea). One or more (5,10,15) diffraction efficiency-compensated holographic spots were sent to the fluorescence sample via the upper microscope objective while z-stack images of each of the light-patterns were acquired by using the inverted microscope and a substage camera and moving the upper objective with a piezoelectric motor (Physik Instruments) within a range of +/- 150 um with 1-µm interval. The axial FWHM of each spot was estimated by fitting the profile to a Lorentzian function.

All the data analysis of optical characterization was done in custom-written script in Python.

### 2P-guided electrophysiological recording *in vivo*

Patch-clamp recordings were obtained under the visual guidance of 2P scanning imaging performed at 920 nm. Positive pressure of >150 mbar was initially applied to the pipette interior filled with the internal solution when the patch pipette penetrated the brain surface and was positioned within or near the opsin-expression region in V1. The pipette pressure was reduced to ∼20 mbar when the pipette tip was close to an opsin-expressing or opsin-non-expressing cell to be patched. Spiking activity of the patched cell was measured by using whole-cell or cell-attached recording in current-clamp configuration to monitor the membrane potential changes. Excitatory current was monitored by performing whole-cell recording in voltage-clamp configuration at -70 mV.

Patch pipette of 5-8 MΩ tip resistance were fabricated from borosilicate glass by using a microelectrode puller (Sutter Instruments). The initial access resistance was 18.2±0.5 MΩ for current-clamp recordings (n=21, opsin-positive and opsin-negative cells in Table 1) and 27.8±3.1 MΩ for voltage-clamp recordings (n=9, opsin-negative cells in Table 1).

The internal solution of patch pipette contained 135 mM K-gluconate, 4 mM KCl, 10 mM HEPES, 10 mM Na_2_-phosphocreatine, 4 mM Mg-ATP, 0.3 mM Na_2_-GTP, and 25-50 µM AlexaFlour-488 for pipette visualization. The external solution, which was applied on top of the craniotomy, contained 145 mM NaCl, 5.4 mM KCl, 10 mM HEPES, 1 mM MgCl_2_, and 1.8 mM CaCl_2_. Membrane potential recordings obtained by using whole-cell current-clamp recordings were corrected for liquid junction potential (11.9±0.2 mV, 5 measurements) offline. Voltage or current recordings were acquired by using a Multiclamp 700B amplifier and a Digidata 1550A digitizer, which were controlled by pCLAMP10 software (Molecular Devices). Electrophysiology data were filtered at 6 kHz and digitized at 20 kHz.

### Quantification of intrinsic electrophysiological properties

Intrinsic electrophysiological properties were accessed by injecting step current-pulse from -150 pA with an increment of 50 pA for 150 ms at current-clamp configuration. The average membrane potential was estimated as the mean membrane potential over 20 ms before current injection. The resting membrane potential was measured as the down-state membrane potential by selecting traces of mean membrane potential that was more hyperpolarized than -50 mV before correcting for liquid junction potential. Input resistance (R_in_) was calculated as the slope of the I-V curve where hyperpolarizing current-pulses were injected (i.e., -150-0 pA) at the down-state ^112^. Membrane time constant (τ_m_) was estimated by fitting an exponential decay for 10 to 95% of the peak hyperpolarization during current injection of -50 or -100 pA. Membrane capacitance (C_m_) was calculated according to τ_m_ = R_in_ * C_m_. Rheobase was determined as the smallest current injection that evoked AP. AP properties were measured according to the first AP evoked at the rheobase. AP threshold was measured as the voltage at which slope exceeded 50 mV/ms. AP amplitude was measured as the voltage between AP threshold and AP peak. AP half-width was estimated as the full-width at half-maximum amplitude. Spontaneous firing rate was accessed as the mean spike rate over 100 ms before current injection and 200 ms after current injection. Evoked firing rate was accessed as the mean spike rate over 150 ms during injection of current that was 100 pA more than the rheobase.

### *In vivo* photostimulation experiment for characterizing presynaptic activation

Single-cell or multi-cell stimulation was performed by using 2P holographic light-shaping with temporal focusing for experiments characterizing the induced AP properties (Fig. 2A-2C) and for experiments characterizing the spatial selectivity of AP generation (Fig. 2D-2G). In a FOV where L2/3 neurons expressed the somatic opsin ST-ChroME, the spiking activity was measured by performing whole-cell current-clamp recordings or cell-attached recordings from the cell soma of a L2/3 neuron expressing the somatic opsin ST-ChroME in V1 of anesthetized mice. The soma coordinates of the patched cell and the surrounding opsin-expressing cells were extracted from 2P scanning imaging and registered manually during experiment. The phase profiles corresponding to holographic light-patterns targeting the patched cell soma (‘1 target’), the 10 surrounding cell somata (’10 targets’), and the 10 surrounding cell somata plus the patched cell soma (’10+1 targets’) were calculated by using fast GPU calculation (∼200 ms/pattern). The phase profiles were to be addressed on the SLM for generating the corresponding holographic light-patterns. According to the diffraction efficiency profile (Fig. 1C, 1D), the stimulation powers required for delivering light-patterns targeting soma at different part of the FOV with a specified power density at the microscope objective (between 0.1-0.5 mW/µm^2^) were calculated. The spiking activity was measured upon brief illumination (between 2-10 ms) of a holographic light-pattern with a specified power density. Stimulation was delivered with an inter-stimulus interval ≥2 s. For measuring the lateral selectivity of AP generation, the light-patterns of single-target or multi-target were delivered at lateral offsets of 0, 10, 20, 30, 50, 70, or 90 µm relative to the patched cell soma by moving horizontally the recording stage where the mouse and the patch recording were situated. For measuring the axial selectivity, the light-patterns of single-target or multi-target were delivered at axial offsets ranging from 0 to ±100 µm above or below the patched cell soma by moving vertically the microscope objective.

### Estimation of off-target activation upon 2P holographic stimulation

We extracted the x-y-z coordinates of all opsin-expressing neurons in 6 representative FOVs by applying automatic cell recognition of Cellpose ^81^ on 2P z-stack red fluorescence images. An additional manual correction allowed to add and remove misidentified ROIs and assured proper identification of all neurons. In each of the FOVs, cells within ± 8 µm from the z=0 central plane were selected as potential target cells (Fig. 2F, red dots) for estimating off-target AP probability.

We then defined a spike probability function SPF as a binary ellipsoid with the lateral and axial size corresponding to the AP FWHM extracted from the AP spatial probability reported in Fig. 2E. If a neighbouring opsin-expressing cell was within the ellipsoid of a target cell, it was considered to be co-activated and contributed to off-target spikes.

The SPF, centered on each of the potential target cells, allowed to estimate if any neighbouring cell was close enough to undergo unwanted photoactivation. The off-target AP probability was then estimated by summing the total off-target activations and dividing it by the number of targeted cells. The estimated off-target AP probability in Fig. 2G was further corrected by multiplying the AP probability at zero spatial offset (≈85%) and the stimulability (≈80%), which was the previously estimated fraction of opsin-expressing cells responding to photostimulation by spiking.

To calculate the cell density reported in Fig. 2F, we divided the number of neurons by the volume of the convex hull that contains all neurons. This approach, with respect to a normalization to the total stack volume, allowed to better estimate cell density under highly localized opsin expression where expressing neurons are clustered and large portions of the stack do not present any opsin-expressing cells.

To obtain the numerical estimation of photostimulation spatial profiles of Fig. S3 we started from the normalized 2P fluorescent profile curve of the holographic spot, lateral *F*_*x*_ and axial *F*_*z*_, reported in Fig. 1B(ii)). Its square root, multiplied by different P values – surface power density values at the central position (either z=0 for axial or averaged around x=0 for lateral) - represent the effective laser light density profiles 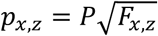. The effect of the power saturation of the opsin responses to different power was then estimated by composing these profiles with a typical 2P response saturation curve of expression 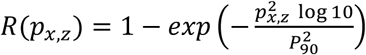, with *P*_90_ the light density to achieve 90% of the maximal response (resulting in the solid profiles in Fig. S3B, C). Additionally, to take into account the size of the cell soma, we performed a convolution of (*p*_*x*,_) with a simplified top hat shape of 15 µm width (resulting in the dashed profiles in Fig. S3B, C).

### *In vivo* photostimulation experiment for connectivity mapping

Single-cell or multi-cell stimulation was performed by using 2P holographic light-shaping with temporal focusing for connectivity mapping experiments. In sequential connectivity mapping, single-cell holographic stimulation was applied for sequentially activating single potential presynaptic cell (Fig. 3). In parallel connectivity mapping, multi-cell holographic stimulation was used for activating sets of multiple potential presynaptic cells and reconstructing individual synaptic responses using CS (Fig. 4, Fig. 5, Table 3).

In a FOV where L2/3 neurons expressed the somatic opsin ST-ChroME, excitatory postsynaptic current was measured by performing 2P-guided voltage-clamp recordings at -70 mV from the cell soma of a L2/3 neuron not expressing opsin in anesthetized mouse V1. Opsin-negative neurons were identified by the absence of the red fluorescence of mRuby, opsin ST-ChroME’s reporter, and a null photoinduced current upon soma illumination (Fig. S2C). The soma coordinates of the patched cell and the surrounding potential presynaptic cells expressing the somatic opsin ST-ChroME were manually identified during experiment. For probing connections from N potential presynaptic cells, the measurement matrices specifying the F cells to be stimulated in each measurement for M measurements were defined. In sequential connectivity mapping (Fig. 3), we designed measurement matrices of F=1 and M=N (diagonal matrix); in parallel connectivity mapping (Fig. 4, 5), we designed measurement matrices of F>1 and M≤N (except FOV5 in Table 3, where M>N). The M phase profiles corresponding to the M light-patterns, which targeted single or multiple potential presynaptic cell somata, were rapidly calculated by GPU (∼200 ms/pattern) and the stimulation powers required for delivering the light-patterns of a specified power density (0.2 or 0.3 mW/ µm^2^) were calculated, taking into account the diffraction efficiency profile and additional losses due to multispot generation (Fig. 1C, 1D). To induce millisecond AP in single or multiple potential presynaptic cells, light-pattern illumination which targeted individual potential presynaptic cell soma and of 10 ms duration and a specified power density of 0.2 or 0.3 mW/µm^2^ was delivered. Single-spot photostimulation patterns were repeated 24-60 times while multi-spot stimulations were repeated 18-60 times. Stimulation patterns were regularly refreshed among repetitions to avoid long and regular stimulation over the same cells. Specifically, in 9 experiments (FOV 1, 2, 5-11 in Table 3), the same stimulation pattern was repeatedly illuminated 6 times at 10 Hz. In 3 other experiments (FOV 3, 4, 12 in Table 3), at each stimulation the holographic pattern was changed (every 100 ms) and the same stimulation pattern was repeatedly illuminated after the completion of a full sequence across the M pattern, requiring Mx100ms. The latter protocol decreased stimulation frequency and allowed therefore minimizing potential synaptic plasticity.

In sequential connectivity mapping, presynaptic cells were directly identified by detecting postsynaptic current responses time-locked to stimulation. In parallel connectivity mapping, connections were retrieved after applying compressive sensing algorithm to the postsynaptic current upon multi-cell stimulation (as better illustrated below).

### Analysis of light-induced AP and photocurrent

Voltage traces were sampled at 20 kHz, filtered for noise of 50 Hz and 150 Hz. The latency of light-induced AP was calculated as the time-span between the illumination onset and the AP peak. The average AP latency was calculated as the mean of AP latencies across 3-8 repetitions and AP jitter as the standard deviation of AP latencies. The AP count was computed as the number of AP induced within 20 ms after illumination onset. The AP probability was computed as the probability of inducing ≥1 AP count. Current traces were sampled at 20 kHz, some filtered for noise of 50 Hz and 150 Hz. The peak current was calculated as the peak of photocurrent within 30 ms after illumination onset. The charge (Fig. S2C) was calculated as the product of current and time within 30 ms after illumination.

### Analysis of postsynaptic currents

Postsynaptic current traces were sampled at 20 kHz, filtered for noise of 50 Hz and 150 Hz, and downsampled to 5 kHz. While each photostimulation pattern targeting single or multiple potential presynaptic neurons was repeated, repetitions of individual postsynaptic current with large fluctuations, i.e., whose amplitudes >2*20 percentile of all fluctuation amplitudes between -20-80 ms relative to the stimulation onset, were first discarded.

Information of the synaptic responses were obtained from the remaining trials, analysing either single repetition traces or average traces across repetitions. Specifically, to identify and evaluate the magnitude of synaptic current responses, we define the quantity time-window average (TWA) as the mean value of a current trace, averaged over a specific time window. TWA is typically computed over interval of few ms before and after presynaptic photostimulation. Focusing primarily on excitatory currents of negative sign, TWAs are often reported, for simplicity, with its sign inverted (positive values corresponding to inward currents), and are defined as TWA amplitudes (TWA amp.).

The individual or average postsynaptic current trace was first baseline subtracted, with a baseline corresponding to the average current in the interval -5 to 0 ms before photostimulation onset; then, we extracted the peak current as the maximum current value within 3 to 25 ms after photostimulation onset.

To declare a synaptic response in postsynaptic current upon single-cell or multi-cell presynaptic stimulation we combined two criteria: 1-we performed a one-sided paired t-test (requiring p<0.05) between the values of TWA of single traces computed in the time interval right before (-5-0 ms) and after (5-20 ms) stimulation; 2-we set a minimum threshold of 2 to 3 pA to the peak current (variable according to the noise of the whole-cell recording (Fig. S6B, S6C)). A target cell was declared as presynaptic connected cell when a postsynaptic current response was identified in sequential connectivity mapping under single-cell presynaptic stimulation. The noise levels of the individual traces for each measurement as well as the noise levels of the traces averaged over repetitions for each measurement were assessed by computing the standard deviation of the current traces within a time interval -5 to 0 ms before photostimulation onset (Fig. S6A-S6C).

The connection rate was calculated for each experiment as the number of presynaptic cells, which were identified in sequential connectivity mapping, divided by the number of all potential presynaptic cells in a FOV.

For individual postsynaptic current traces that were declared as responses, we defined the response onset values, computed for individual traces as TWA of baseline current between -5-0 ms (from stimulation onset) minus 2 standard deviations of the baseline current in the same interval. No responses were identified as individual traces where TWA between 10-15 ms did not reach their onsets. Visual inspection was then applied for validating responses or failures in individual traces. The response rate of a connection was determined as the ratio between repetition number of no failure traces and the number of non-discarded repetitions.

As for the kinetics of the average postsynaptic current traces, similar to before, response onsets were computed as the TWA in the interval -5-0 ms minus 2 standard deviations of the average current traces in the same interval. Latencies were analysed as the time in which the average current reaches a negative value corresponding to its response onsets (restricted to the interval 0-15 ms from the stimulation onset). Peaks were identified as the local minima of the average traces within 5-25 ms from the stimulation onset. Peak latencies were analysed as the time of the peaks, and peak amplitudes the differences between onsets and peaks. Rise times were calculated as the time differences between 20% and 80% peak amplitudes. Offsets were identified as the time reaching the values of onsets after the response peak. Half-widths were calculated as the time span between the rising and decaying phases of responses reaching 50% peak amplitudes. Decay time constants were calculated by fitting exponentials from peaks to offsets by using the least-squares method and of an upper bound of 100 ms.

To compare the current responses upon multi-cell stimulation vs the sum of those upon single-cell stimulation (Fig. 4) and to reconstruct synaptic weights by combining CS and responses upon multi-cell stimulation (Fig. 5, Fig. S6D), TWA amplitudes of the averaged current traces after pre-processing (filtering, down-sampling, discarding traces of large fluctuation, and baseline subtraction) were computed. Of note, TWA amplitudes corresponding to excitatory currents are reported as positive values, and since we focused on the excitatory synaptic inputs, negative values of TWA amplitudes (corresponding to outward currents) were offset to zero. The choice of the time interval for computing the TWA, here 10-15 ms after stimulation onset, was related to the previous characterization of peak latency under single-cell stimulation (Fig. 3C), according to which most of current response peaks were included in this time interval; nevertheless, the analysis performed with other time interval (i.e. 5-20 ms after stimulation onset) didn’t significantly affect the results of input summation (results not shown).

### Effect of off-target activation on estimated connection rate

The prediction of the effect of off-target stimulation on the estimated connectivity rate (Fig. S4) were based on the following expression for the probability of detection of a synaptic response under single-spot stimulation:

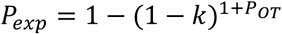

where *k* is the true connectivity rate and *P*_0*T*_ is the probability of inducing off-target APs. The expression, also present in ^70^, shows how, with *P_OT_* > 0, *P*_*exp*_ > *k*, resulting in a potential overestimation of the connectivity rate. In the figure, the experimentally estimated connectivity rate, and its absolute *P*_*exp*_ − *k* and relative (*P*_*exp*_ − *k*)⁄*k* error are plotted as a function of *k* and *P_OT_*. For low values of off-target probability, the expression can be developed to the first-order giving : *P*_*exp*_ ≈ *k* − (1 − *k*) · log(1 − *k*) · *P_OT_* = *k* + *c* · *P_OT_*., with *c* > 1. In Fig. S4B this expression is plotted showing how, for low *P_OT_*, the overestimation in connectivity rate is directly proportional to the off-target probability.

### Experimental compressive sensing reconstruction of connectivity

Compressive sensing aims to find solutions to an underdetermined problem under incoherent probing of a system that is sparse and linear ^76,77^.

In parallel connectivity mapping, to probe the connectivity to the postsynaptic cell from N potential presynaptic cells (x _N×1_), M measurements of postsynaptic currents upon stimulating subsets of multiple potential presynaptic cells according to the measurement matrix (A _M×N_) were recorded ^72^. Incoherent probing was achieved by defining a stimulation matrix in which each row contains F non-zero elements corresponding to the targeted neurons in a photostimulation pattern, where the N cells were distributed randomly over the M stimulation patterns, with the only constraint of being sampled as homogenously as possible (each cell stimulated a similar number of times).

As the feature input of the CS reconstruction, we considered the TWA amplitudes of the average current traces induced by each multi-cell stimulation (calculated in the interval between 10-15 ms after stimulation onset). These scalar values defined the measurement vector y _Mx1_. The goal of CS reconstruction was therefore to solve for x in *y*_*M*×1_ = *A*_*M*×*N*_*x*_*N*×1_. To achieve it, we applied the basis pursuit solution to minimize over x the loss function

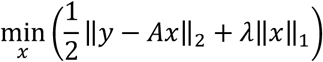

where λ was a regularization hyperparameter which influenced the sparsity of the solution and prevented overfitting. This was implemented by using a CVX Matlab package ^113,114^ for convex optimization models, similar to ^74^. In the present study, λ=0.1 was applied for connectivity reconstruction. The N output values of the reconstructed algorithm correspond to the TWA amplitudes of the postsynaptic currents upon single-cell stimulation of N potential presynaptic cells, which represent the synaptic strength of connections.

To define connected or non-connected cell from the reconstruction output, k-means clustering for 2 clusters was applied to the reconstructed connectivity vector x, one cluster defining retrieved connections and the other defining non-connected cells (as shown in Fig. 5A(ii, iii), 5B(ii, iii)).

The performance of compressive sensing in parallel connectivity mapping was evaluated by comparing the reconstruction result with the ground truth connectivity determined in sequential connectivity mapping. More precisely, true positives (TP), true negatives (TN), false positives (FP), and false negatives (FN) were determined by verifying whether reconstructed connected or non-connected cells corresponded to the identification obtained under single-cell sequential mapping (discussed and reported in Fig. 3, Table 3). Precision was computed as TP/(TP+FP), indicating the relevance of all reconstructed connections. Recall was computed as TP/(TP+FN), specifying the retrieval of all reconstructed connections. Receiver operating characteristic (ROC) curves were computed to visualize the performance of the CS reconstruction (Fig. S12A), in terms of true positive rate (y-axis) and false positive rate (x-axis) at all classification thresholds. The area-under-curve (AUC) of the ROC curves was then computed as one of the evaluation metrics.

To examine how different parameters, including the number of measurements *m*, regularization parameter λ and connection-defining threshold, affected the performance of CS reconstruction of network connectivity, reconstruction was performed by under a grid search of these parameters (Fig. S12 B-F). Subsampling measurement number *m* was smaller or equal to the total measurement number. The corresponding compression ratio (CR), which indicates the speed gain, was computed by CR=N/m. For each subsampling value (excluding m=M), different subsets of *m* measurements out of the total number of measurements were randomly chosen, and the number of subsets was equal to M in order to balance sampling across all potential presynaptic neurons, meaning that in each subsets each potential presynaptic neuron were sampled by similar number of measurements. Mean and S.E.M. of the reconstruction responses and performances for each of the subsampling value were obtained by averaging the value obtained across these subsets of measurements.

### Simulation of compressive sensing connectivity reconstruction

Similar to what has been done in ^74^, we considered an observed network of either N=30 Izhikevich model neurons with an average of K=2 connections (≈6.7% connectivity) or N=100 neurons with an average of K=10 connections (10 % connectivity) within a 1000 neuron network. The Izhikevich model was chosen for its good balance between biological plausibility, diversity of firing characteristics and computational simplicity. A ratio of 80% excitatory positive postsynaptic weight and 20% inhibitory negative post-synaptic weight was chosen to mimic mammalian cortex. Neurons were connected with uniform random probability. In order to produce the desired level of spontaneous firing, each neuron was independently excited with different current sampled from a zero mean gaussian distribution. The variance of the current was tuned to evoke a firing rate of 0.2 Hz per neuron averaged across the network. These parameters were chosen based on reported spontaneous activity in layer 2/3 V1 in awake mice ^70,106,115,116^. We generated an M x N binary random stimulation matrix A where each took the value of ‘1’ with probability F/N and ‘0’ otherwise. For each measurement, a row was randomly selected without replacement from A. Neurons corresponding to the indices in A were forced to fire and the induced postsynaptic current on one specific neuron was monitored. Specifically, peak current in the time interval [0-5 ms] after the synchronized stimulation-evoked presynaptic spikes defined the features vector *y*, input for basis pursuits reconstruction. Values of the resulting vector *x* which were at least as strong as 2% of the maximum possible weight in each network configuration, were considered connections and used to evaluate precision, recall and accuracy metrics. Results reported in Fig. S10 are obtained varying over different values of M and with F varying between 5 and 30% of N and each condition is simulated over 500 different network configurations.

### Statistical analysis

Statistical tests were carried out by using Matlab software. Data comparisons between experimental conditions (e.g., optical properties of holographic light-patterns of different spot numbers, AP properties upon stimulation of different holographic light-patterns) were performed by using ANOVA in combination with multiple comparisons of Tukey’s method, paired t-test (e.g., TWA of individual postsynaptic current traces before and after presynaptic photostimulation), Wilcoxon signed rank test (e.g., off-target AP probability upon stimulation of ‘1 target’ and ‘10+1 targets’), or two-sample t-test (e.g., electrophysiological properties of opsin-positive vs opsin-negative neurons). Data was presented as mean±s.e.m. if not otherwise indicated. Error bars were represented s.e.m. if not otherwise indicated.

## Supplementary Figures and Tables

**Figure S1:**
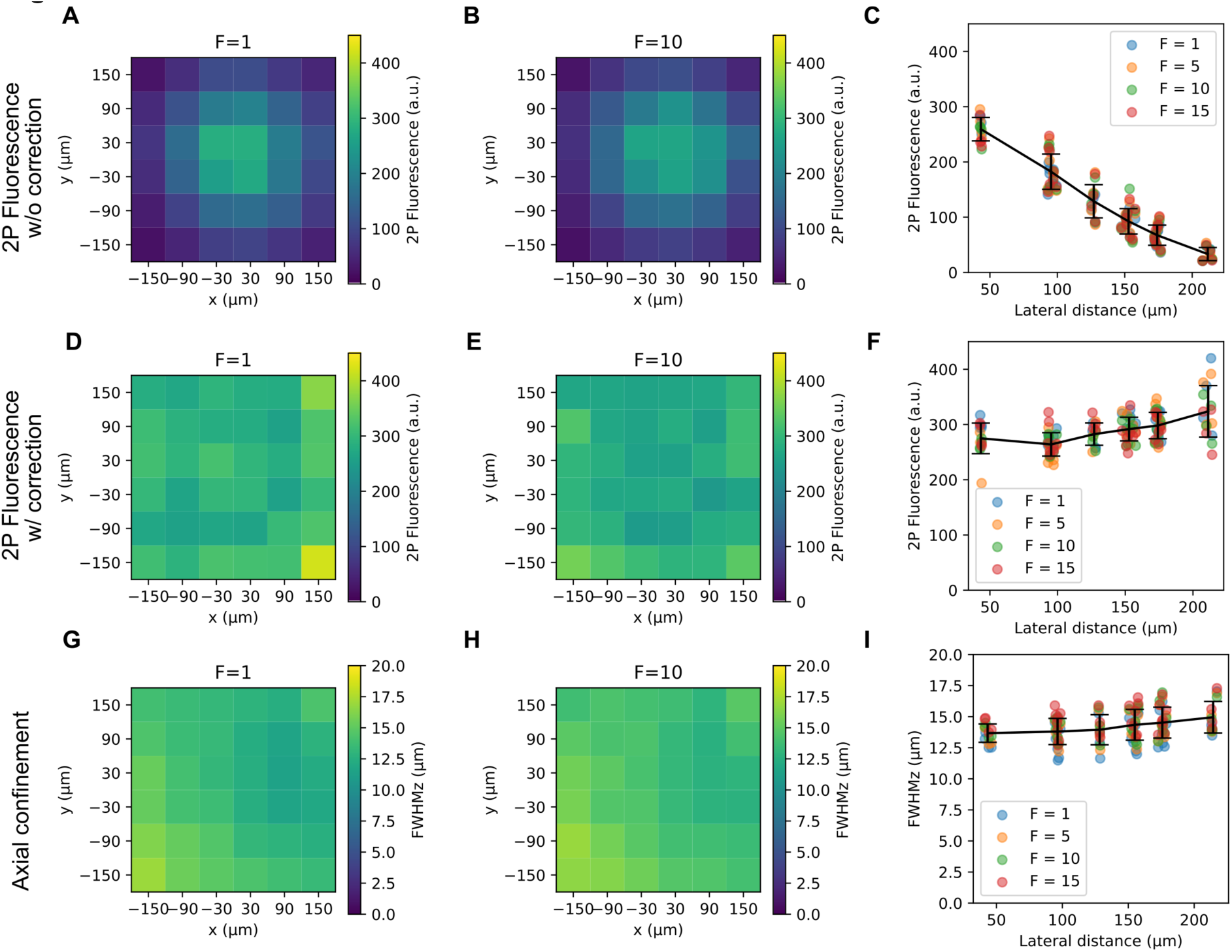
Spatial dependence of the optical properties of the photostimulation spots of the dataset in Figure 1D. Multiple holograms were generated containing F=1, 5, 10 or 15 light-spot(s) each, randomly placed in a 6-by-6 grid with small random offset within a FOV of 300 µm x 300 µm (same dataset as Figure 1C, 1D). (A-C): Spatial distribution of 2P fluorescence of light-spots as a function of their x-y positions (A for F=1 spot per hologram, B for F=10 spots per hologram) or as a function of their distance from the center of the FOV (C), obtained without diffraction efficiency correction. (D-F): Similar to A-C, obtained with correction of diffraction efficiency. In panel A-F, data points for each spot position were averaged over 1 to 6 holograms from a total of 222 holograms (n=36, 40, 20 and 15 holograms each generating 1,5,10,15 spot(s) respectively, separate datasets with and without diffraction efficiency correction). (G-I): Similar plots of A-C, representing the spatial distribution of axial confinement (FWHMz) of holographic spots. In panel G-I, data points for each spot position were averaged over 1 to 3 holograms from a total of 74 holograms (n=36, 20, 10 and 8 holograms each generating 1,5,10,15 spot(s) respectively). Error bars as mean±s.d.

**Figure S2:**
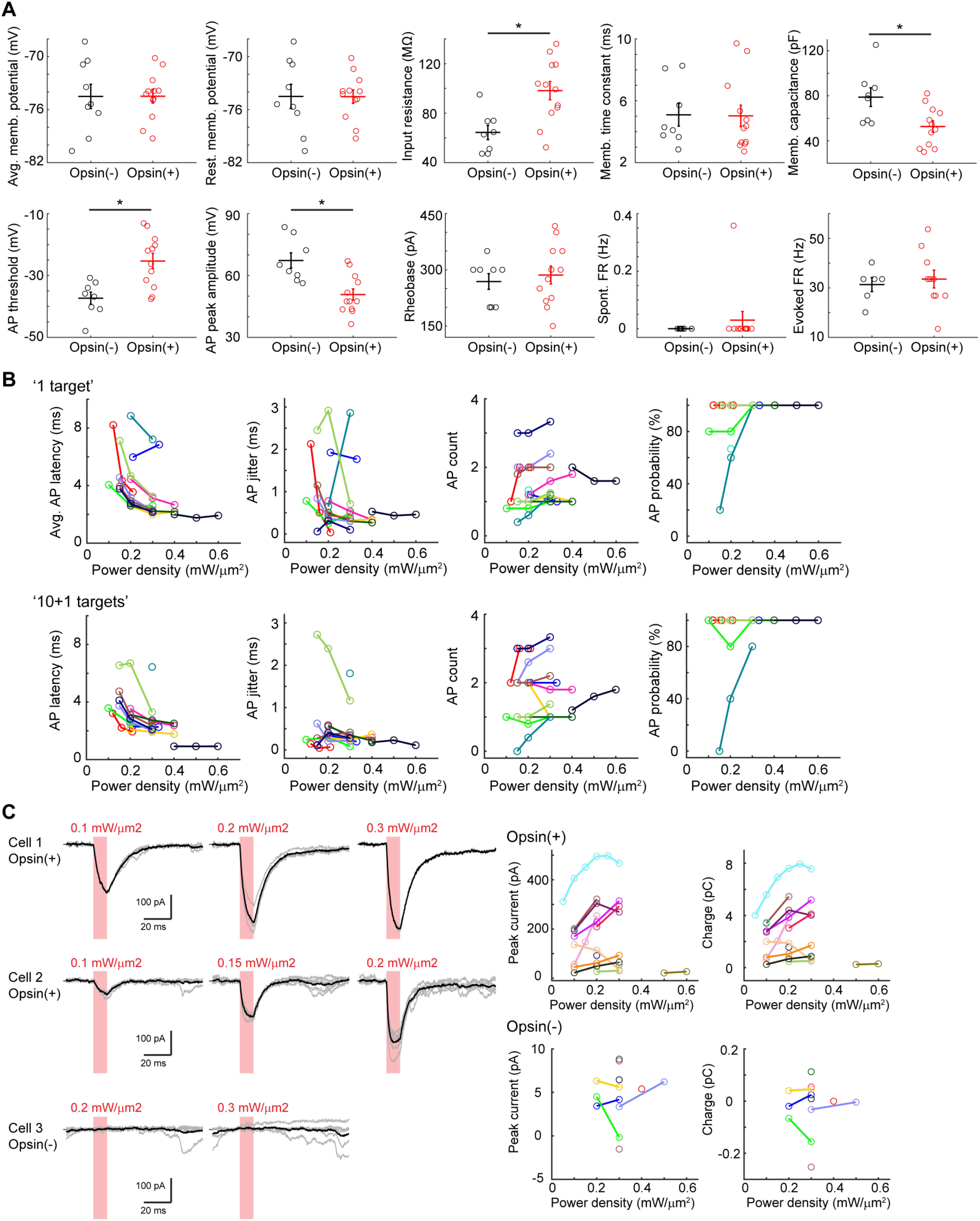
In vivo measurements of 2P holographic stimulation of L2/3 neurons expressing somatic ST-ChroME opsins at anesthetized mouse visual cortex, related to Table 1 and Figure 2A-2C. (A) Electrophysiological properties of opsin-positive (Opsin(+)) and opsin-negative (Opsin(-)) cells, related to Table 1. Opsin(-) cells were denoted in black and Opsin(+) cells were denoted in red (n=9, 12 for Opsin(-), Opsin(+) cells). Opsin(-) cells here were the postsynaptic cells in connectivity experiments. Significant difference in electrophysiological properties between Opsin(-) and Opsin(+) cells denoted as asterisk (two-sample t-test, p<0.05). (B) AP properties as functions of stimulation power density. AP were induced upon 10-ms illumination, related to Figure 2A-2C. Upper, AP properties induced upon stimulation of ‘1 target’ (n=12, 12, 13, 13 for average AP latency, AP jitter, AP count, AP probability). Lower, AP properties induced upon stimulation of ‘10+1 targets’ (n=12, 12, 12, 12 for average AP latency, AP jitter, AP count, AP probability). Data of the same cell were denoted in the same color in both upper and lower panels. (C) Photocurrent properties as functions of stimulation power density. Photocurrent was induced upon single-cell stimulation of 10-ms illumination duration. Left, current traces upon photostimulation of varying stimulation power density in three example cells. Cell 1 and Cell 2 were Opsin(+) and Cell 3 Opsin(-). Individual current traces denoted in grey and average current traces in black. Right, peak current and charge of average photocurrent upon photostimulation of varying stimulation power density. Data of the same cell were denoted in the same color in both upper and lower panels (n=14, 9 for Opsin(+), Opsin(-) cells).

**Table S1:** AP properties upon single-cell, 10-ms 2P holographic stimulation of different power density ranges1.

| | <0.15 mW/$\mu\text{m}^2$ | 0.15-0.3 mW/$\mu\text{m}^2$ | >0.3 mW/$\mu\text{m}^2$ |
| --- | --- | --- | --- |
| AP latency (ms) | 5.85 ± 0.72 (n=6) | 5.09 ± 0.38 (n=29) | 4.76 ± 0.67 (n=14) |
| AP jitter (ms) | 0.97 ± 0.31 (n=6) | 0.99 ± 0.14 (n=29) | 0.83 ± 0.18 (n=14) |
| AP count | 0.77 ± 0.23 (n=11) | 1.08 ± 0.11 (n=37) | 1.15 ± 0.10 (n=17) |
| AP probability (%) | 58.79 ± 14.41 (n=11) | 81.13 ± 5.34 (n=37) | 92.16 ± 5.37 (n=11) |

Data were presented as mean ± s.e.m.

**Figure S3:**
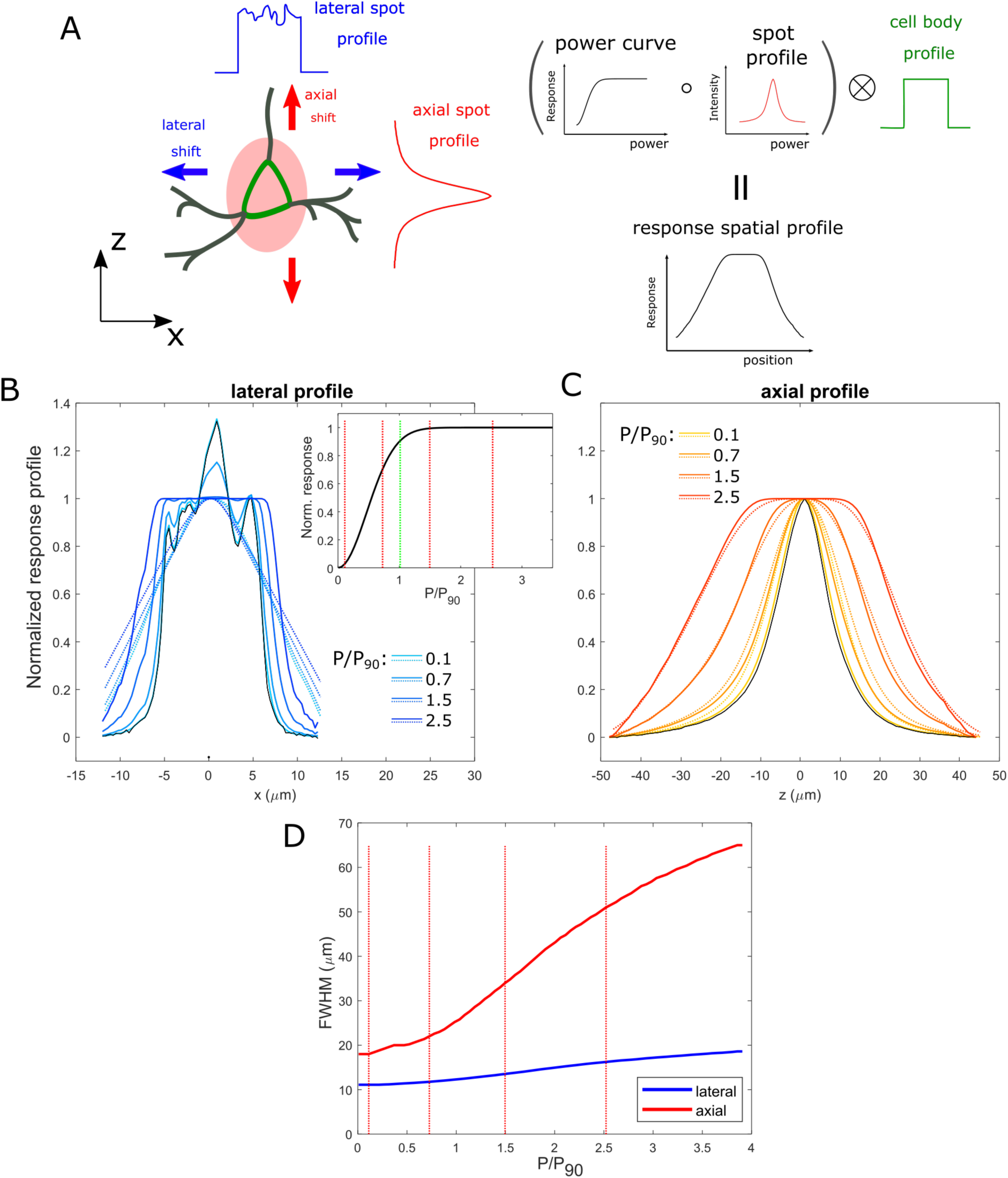
Simulated effect of photostimulation saturation on lateral and axial profile of photocurrents. (A) Schematic of the approach to numerically evaluate the effect of different light density on the photostimulation spatial profiles (lateral and axial), based on the combination between the measured optical holographic PSF and the opsin response curve at different powers and the convolution with the size of the target cell body (see Methods for details). (B) Lateral holographic spot fluorescence profiles (black, optical PSF from profiles of Figure 1B) and corresponding simulated lateral profiles of the photostimulation response at increasing stimulation power P = 0.1, 0.7, 1.5 and 2.5 * P_90_ (light to dark blue solid line) obtained using a typical photocurrent versus illumination power curve shown in the inset, where the dashed green line indicates the light intensity value P_90_ (the value inducing 90% of the maximal response) and the dashed red lines indicate the values corresponding to the profiles of the main panel (P=0.1, 0.5, 1 and 2.5 * P_90_). Dotted lines (from light to dark blue) in main panel represent the response profile obtained after the additional convolution with a 15-µm top-hat profile, representing the extension of the cell soma. (C) Same as B for the axial profile: in solid black the optical axial profile, in solid from orange to red the simulated photoresponses profiles at increasing stimulation power, in dashed from orange to red the corresponding curve after convolution with the cell profile. (D) FWHM of the lateral (blue) and axial (red) dotted profiles in A and B, as a function of the illumination intensity P/P_90_.

**Figure S4:**
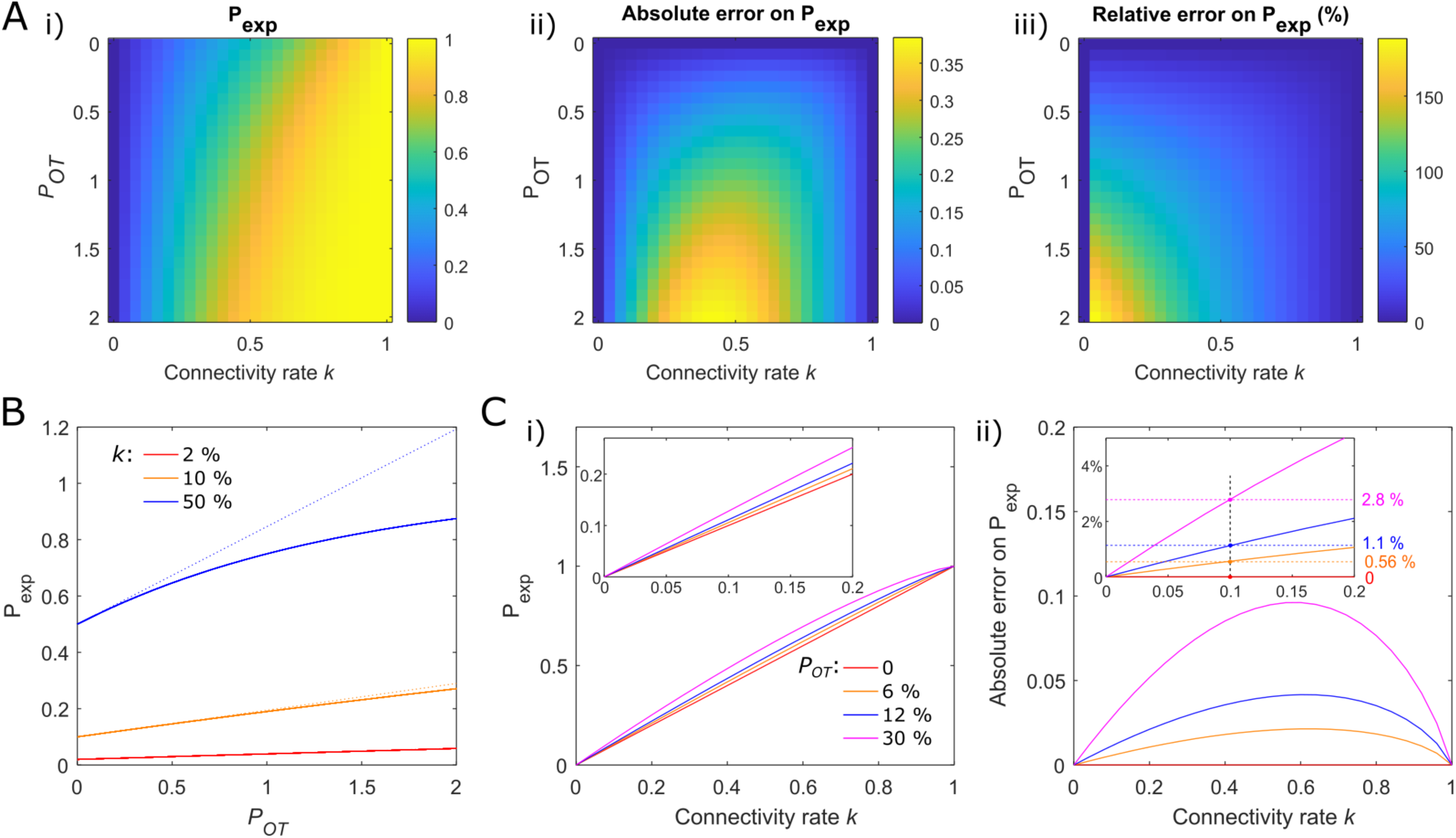
Effect of off-target stimulation on the estimation of connectivity rate. (A) Colormap representing: (i), the experimentally estimated connectivity rate *P_exp_* (i.e. the probability of detecting a synaptic responses upon single-cell sequential stimulation), (ii), the corresponding absolute evaluation error (*k-P_exp_*) and (iii) the relative error (*(k-P_exp_)/k* as a function of the true connectivity rate *k* and the off-target AP probability *P_OT_*, following the expression *P*_*exp*_ = 1 − (1 − *k*)^1+*P*_*OT*_^ (see Methods). (B) Vertical profiles of the colormap in A(i) showing the response probability *P_exp_* as a function of off-target probability *P_OT_* for different representative values of nominal connectivity rate *k* of 2%, 10% and 50%. Dashed lines represent the first order expansion of *P_exp_* expression for low *P_OT_* values. (C) Estimated connectivity rate (i) and its absolute error(ii) as a function of nominal connectivity rate *k* for different values of the off-target probability *P_OT_* : 0, 6, 12 and 30%. The perfect diagonal curve for *P_OT_* =0, corresponds to the correct predictions *P_exp_=k*. In the insets, a zoom in the experimentally relevant region of sparse connectivity *k*=0-0.2. The vertical black dashed line indicates connectivity rate of 10% and the intersects with the colored curve indicates the corresponding absolute error on the estimated connectivity rate (reported on the right) for the different values of *P_OT_*.

**Figure S5:**
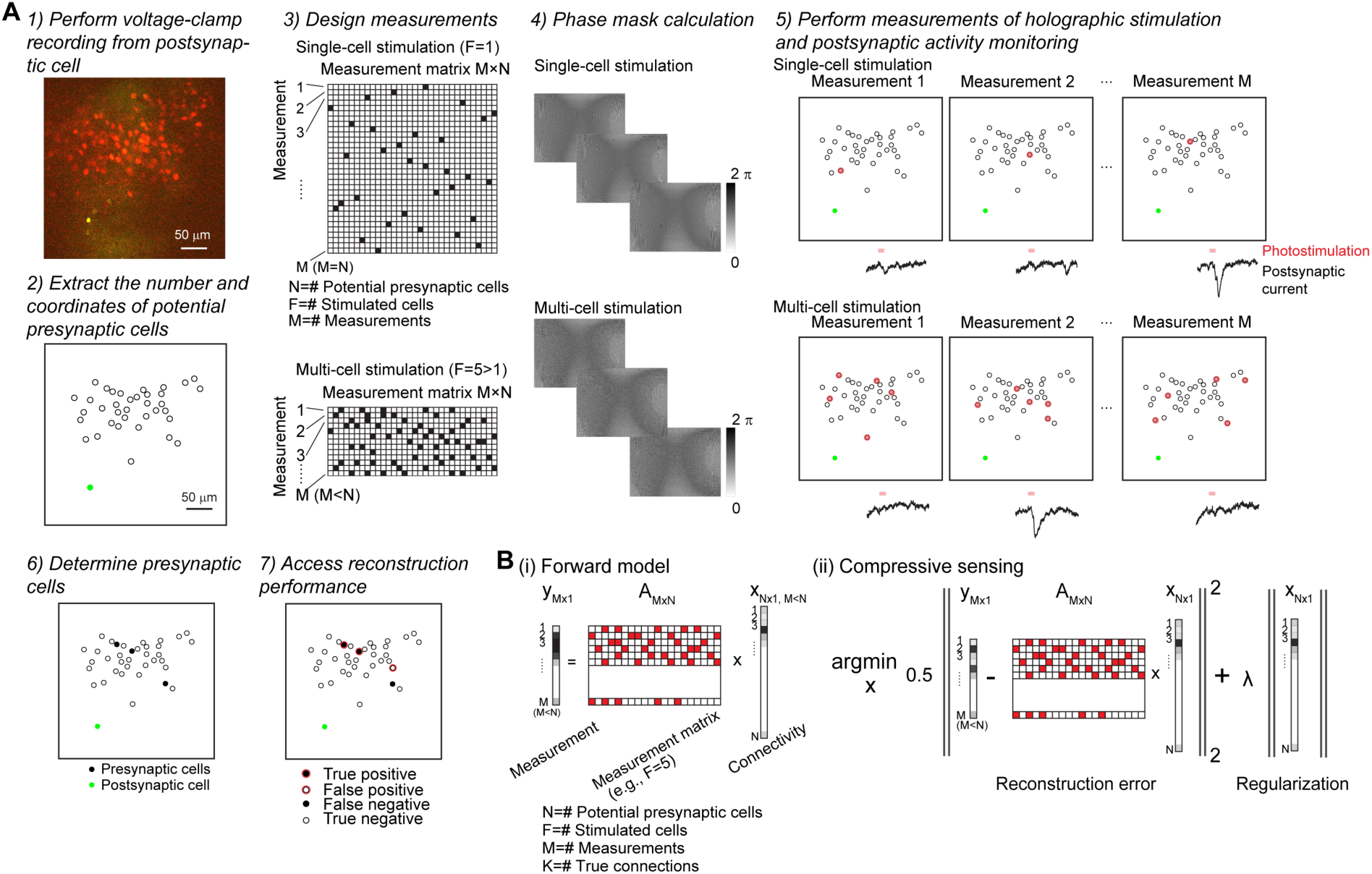
Pipeline of *in vivo* connectivity mapping experiments. (A) Experimental pipeline. Step 1) The excitatory current activity was recorded by using voltage-clamp recording at -70 mV in an opsin-negative postsynaptic cell at L2/3 of anesthetized mouse V1 appearing as a green cell due to the diffusion of the Alexa-488 from the patch pipette. A 2P image of a FOV for an example experiment is shown. Potential presynaptic cells expressing the somatic opsins ST-ChroME appear as red cells in FOV. Step 2) The coordinates of the potential presynaptic cells and the postsynaptic cell were extracted manually or by using a software for automated cell detection. In the example FOV, potential presynaptic cells illustrated as black open circles, and the postsynaptic cell as green filled circle. Step 3), the measurement matrices of single-cell stimulation and multi-cell stimulation, taking into account the parameters of N, M, F, were generated for performing sequential connectivity mapping and parallel connectivity mapping respectively. Step 4), the corresponding phase masks for generating holographic light-patterns targeting single cell or multiple cells were computed by using Gerchberg-Saxton algorithm implemented under GPU calculation. Step 5) Photostimulation under single-cell stimulation or multi-cell stimulation was performed. The light-power to be delivered for each light-pattern was calculated, taking into account the inhomogeneity of diffraction efficiency across the FOV and additional power losses due to multisport generation. During photostimulation, the postsynaptic current activity was monitored. Target cells illustrated as red shaded circles. Step 6) The presynaptic cells connecting to the postsynaptic cell were identified according to the postsynaptic current upon single-cell stimulation. Presynaptic cells illustrated as black shaded circles. Step 7) The connectivity was reconstructed by applying compressive sensing to postsynaptic activity upon measurements of multi-cell stimulation. The reconstructed connections were illustrated as red open circles. True positives, false positives, false negatives, and true negatives were identified by comparing the ground-truth connections in black shaded circles and the reconstructed connections in red open circles. (B) Illustration of the forward model and compressive sensing reconstruction. (i): Forward model of a sparse and linear system which approximate the synaptic integration at the postsynaptic cell upon a sequence of multi-cell stimulation defined by measurement matrix A. (ii): Cost function of compressive sensing, consists of an error term and a regularization term, which was minimized to reconstruct the connectivity vector x.

**Figure S6:**
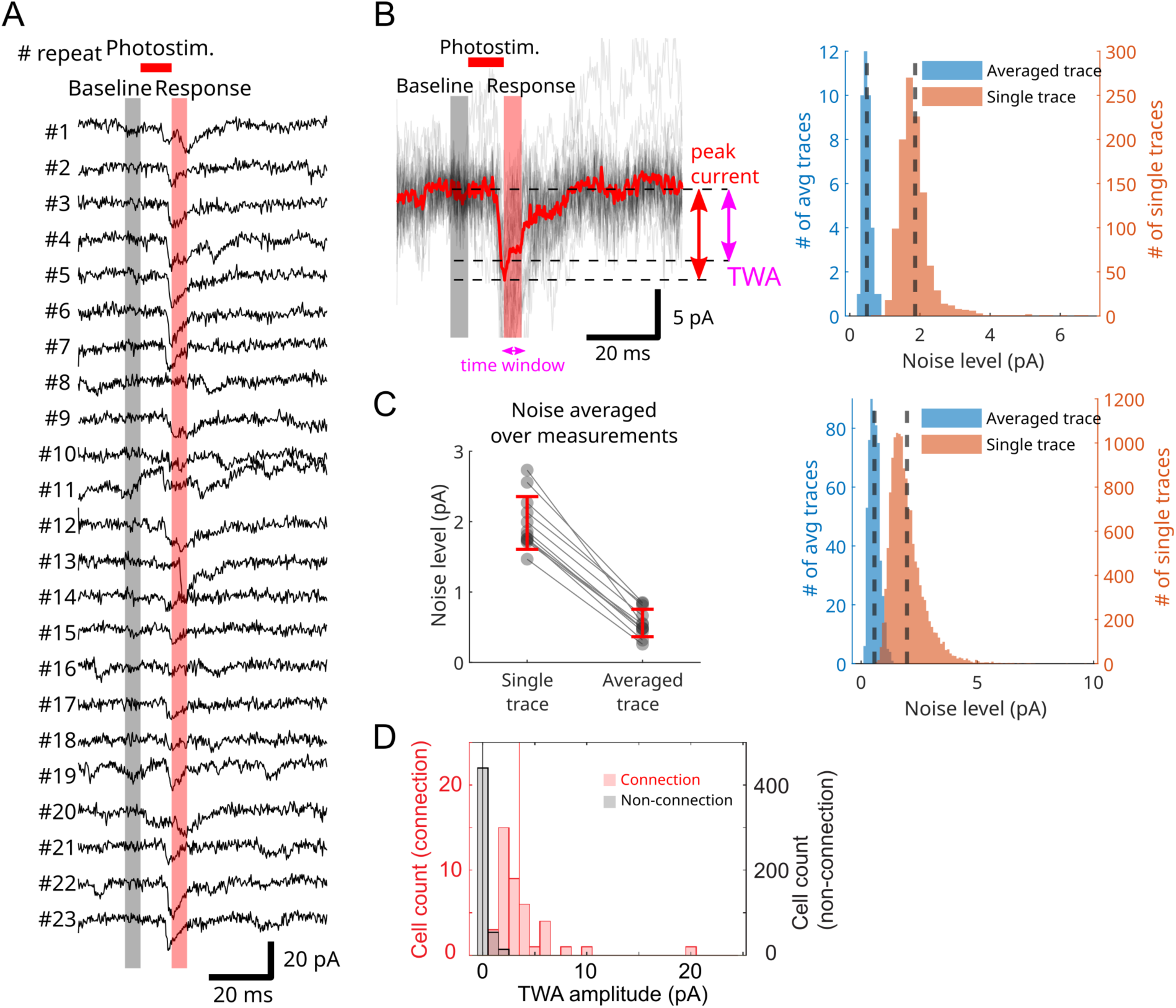
Post-synaptic currents and noise from postsynaptic cells recording. (A) Example of individual traces upon repeated stimulation of a connected presynaptic cell. Baseline as 5 ms before photostimulation of presynaptic cell. Photostimulation occurred between 0-10 ms. Responses were measured 10-15 ms after onset of photostimulation. (B) Left: Superposition of curves in A. Average trace across 23 repetitions is indicated in red. To identify and evaluate magnitude of responses, from average trace we extracted either the peak current (maximum of averaged current after photostimulation event, red arrow) or the time-window average (TWA, purple arrow), meaning the mean current value of the average current trace in a define time window (here from 10 to 15 ms from photostimulation onset). Right: Histograms of baseline noise level of the single and averaged traces of an experiment (1404 repetitions from stimulation of 39 presynaptic cells in 1 example FOV). Vertical dotted gray lines indicate the average value of the respective histogram. (C) Left: Average noise level of the single and averaged traces across 12 experiments (mean±s.d.). Right: Histogram of baseline noise level (20598 repetitions from stimulation of 549 measurements in 12 FOVs). Vertical dotted gray lines indicate the average value of the respective histogram. (D) Histograms of the TWA amplitudes between 10-15 ms after photostimulation onset for the averaged postsynaptic current traces upon stimulation of single cells identified as connections (red, n=41 cells, 12 experiments) or non-connections (black, n=549 cells, 12 experiments). Vertical colored lines denoted medians.

**Figure S7:**
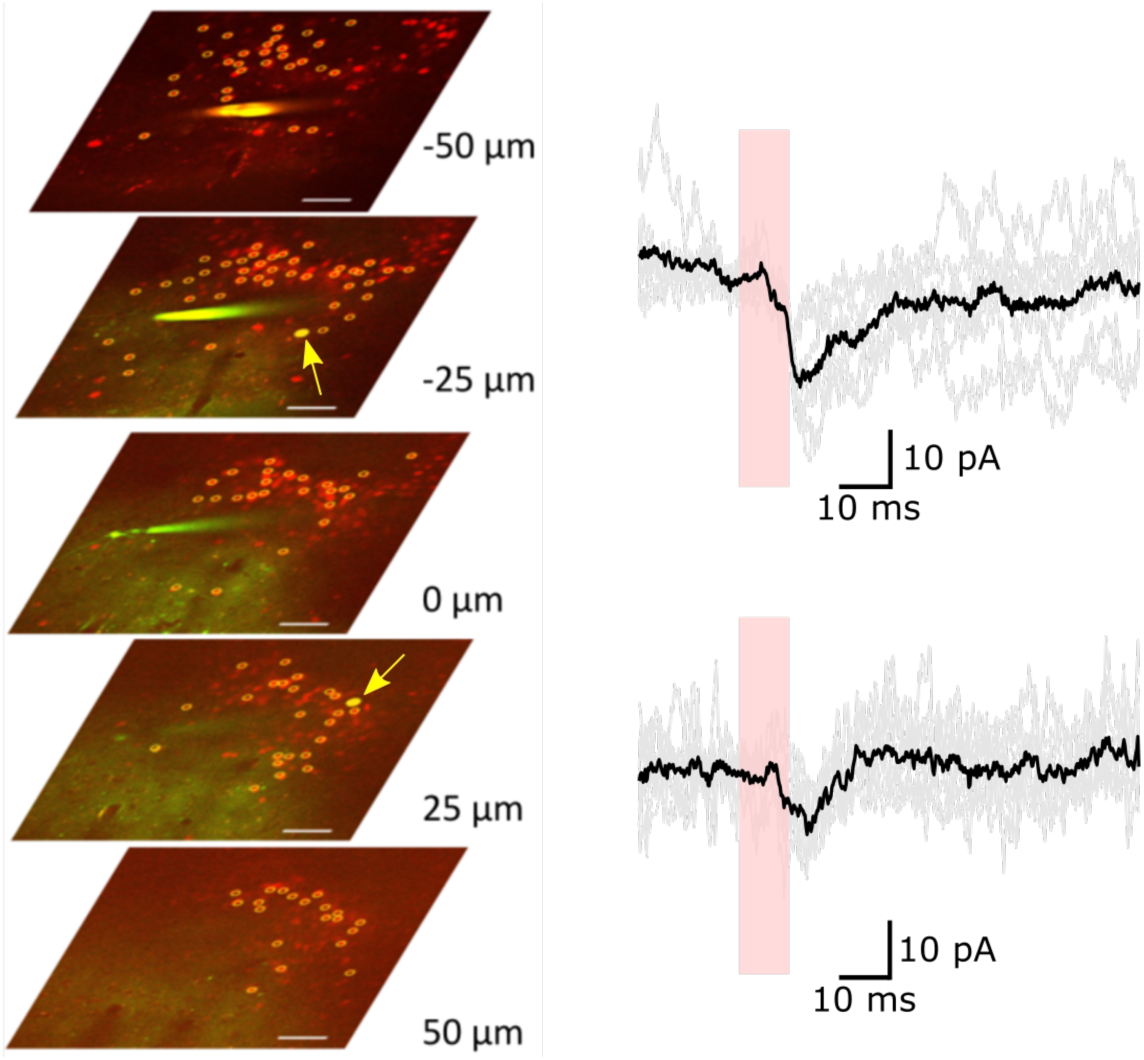
Connectivity mapping in 3D by using single-cell stimulation. Sequential mapping across 137 potential presynaptic cells distributed at 5 different axial planes and expressing the somatic opsins ST-ChroME (red cells in the 2P images denoted with yellow circles), while monitoring excitatory postsynaptic current of a patch cell located at z=0 (corresponding to the depth of 106 µm in the tissue). The patch pipette was visible, by the green fluorescence of Alexa-488 in the internal solution. Current traces on the right represent examples of responses from the stimulation of 2 identified presynaptic cells, indicated by yellow filled circles and arrows on the 2P images, located at the depth of +25 and -25 µm with respect to the patch cell. Illumination conditions: 0.2 mW/µm^2^, 10 ms pulses, n=7, 9 repetitions. Scale bar 50 µm.

**Figure S8:**
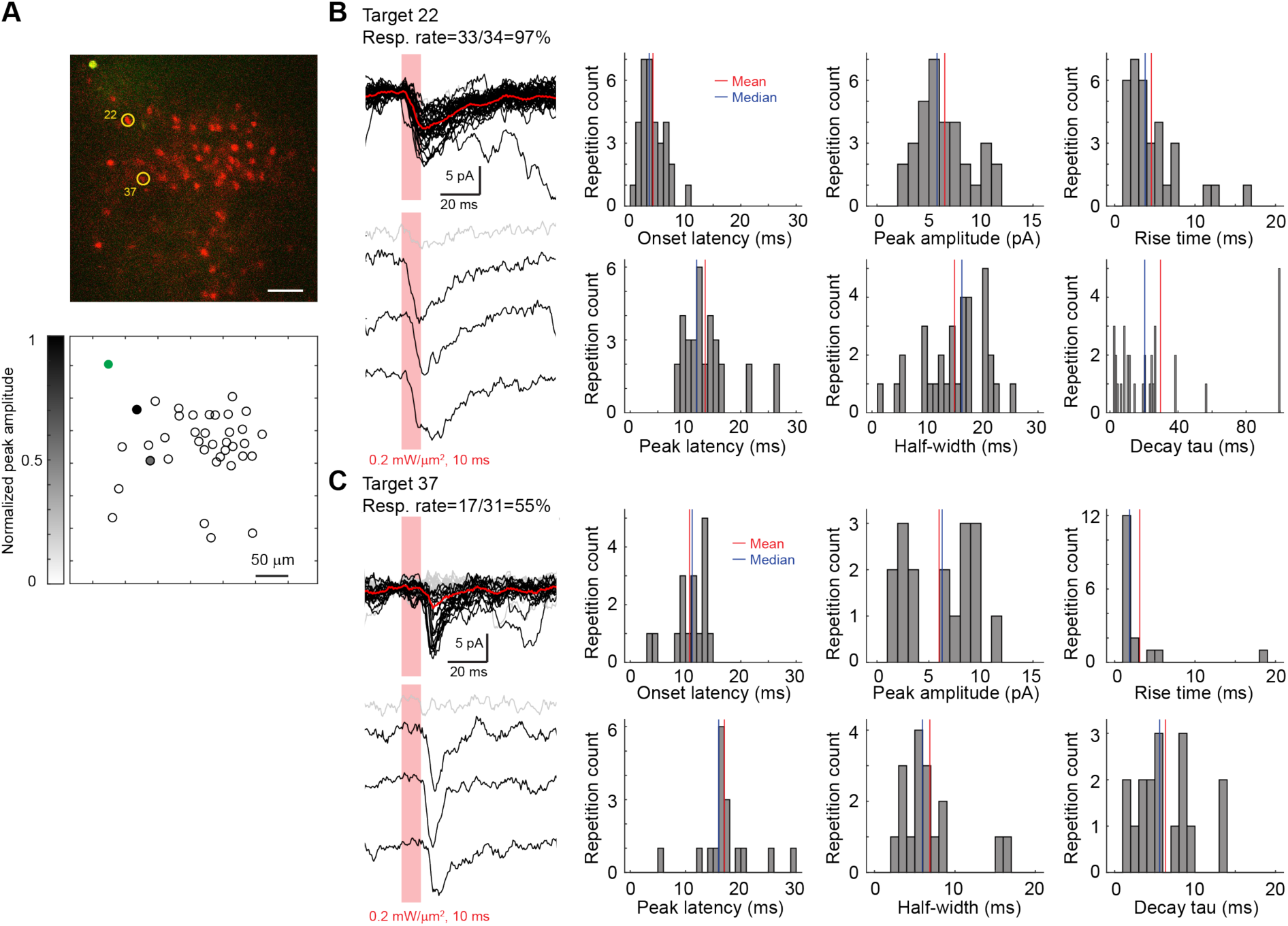
Response kinetics of individual current traces for two exemplary connections identified by using single-cell stimulation in one FOV, related to Figure 3. (A) Example experiment of connectivity mapping by using single-cell stimulation, related to Figure 3A. Upper, 2P image of the opsin-negative postsynaptic cell (green cell soma at the upper left of FOV) and the opsin-positive potential presynaptic cells (red cell soma indicating expression of somatic opsin ST-ChroME). Two identified presynaptic cells (Target 22, Target 37) denoted as yellow circles. Lower, cell coordinate map of the FOV. The postsynaptic cell indicated in green, the potential presynaptic cells as black open circles, and the 2 identified presynaptic cells as grey circles whose shade color denoting the normalized postsynaptic response amplitudes. (B) Individual traces of postsynaptic current responses upon single-cell stimulation of Target 22. Upper left, overlaid postsynaptic current traces upon 34 repetitions of presynaptic stimulation of Target 22, in which 33 responding traces are indicated in black and 1 not responding trace in grey. The average trace across 34 repetitions denoted in red. Lower left, example postsynaptic current traces upon 4 repetitions of presynaptic stimulation. Photostimulation denoted as red shades. Right, histograms of kinetics and amplitude of responding postsynaptic current traces. (C) Individual traces of postsynaptic current responses upon single-cell stimulation of Target 37. Upper left, overlaid postsynaptic current traces upon 31 repetitions of presynaptic stimulation of Target 37, in which 17 responding traces indicated in black and 14 not responding traces in grey. The average trace across 31 repetitions denoted in red. Lower left, example postsynaptic current traces upon 4 repetitions of presynaptic stimulation. Photostimulation denoted as red shades. Right, histograms of kinetics and amplitude of responding postsynaptic current traces.

**Figure S9:**
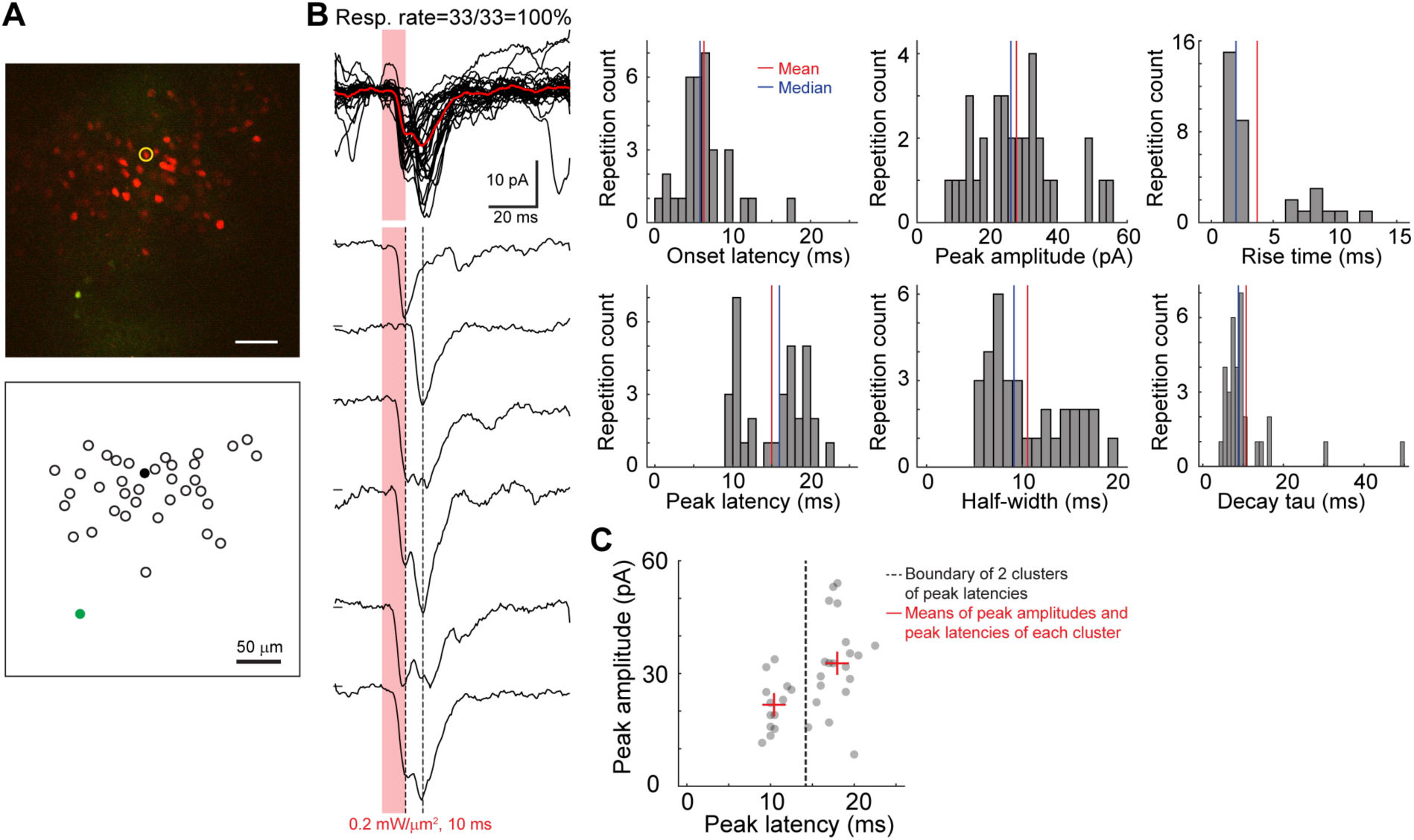
Response kinetics of individual current traces for an exemplary connection identified by using single-cell stimulation in one FOV. (A) Example experiment of connectivity mapping by using single-cell stimulation. Upper, 2P image of the opsin-negative postsynaptic cell (green cell soma at the lower left of FOV) and the opsin-positive potential presynaptic cells (red cell soma indicating expression of somatic opsin ST-ChroME). One identified presynaptic cell denoted as yellow circle. Lower, cell coordinate map of the FOV. The postsynaptic cell indicated in green, the potential presynaptic cells as black open circles, and the 1 identified presynaptic cell as black shaded circle. (B) Individual traces of postsynaptic current responses upon single-cell stimulation of the identified presynaptic cell. Upper left, overlaid postsynaptic current traces upon 33 repetitions of presynaptic stimulation. Lower left, postsynaptic current traces upon 6 repetitions of presynaptic stimulation. Postsynaptic current traces displayed peaks of varying latencies, which were denoted by the 2 dashed lines. Photostimulation denoted as red shades. Right, histograms of kinetics and amplitude of responding postsynaptic current traces. (C) Peak amplitudes vs. peak latencies of responding postsynaptic traces. Peak latencies were classified in 2 clusters by using k-means clustering. The correlation coefficient between peak latencies and peak amplitudes was 0.49.

**Table S2:** Kinetics of average response traces upon single-cell stimulation of presynaptic partner neuron.

| Onset latency (ms) | Peak latency (ms) | Peak amplitude (ms) | Rise time (ms) | Decay time constant (ms) | Half-width (ms) |
| --- | --- | --- | --- | --- | --- |
| 5.8±0.4<br>(n=41 connections) | 14.6±0.4<br>(n=41) | 5.0±0.6<br>(n=41) | 4.3±0.3<br>(n=41) | 10.7±1.0<br>(n=35)* | 11.8±0.9<br>(n=41) |

\* Calculated for decay time constants < 30 ms; 16.4±2.6 ms for decay time constants of all 41 connections in 12 FOV
Data were presented as mean±s.e.m.

**Figure S10:**
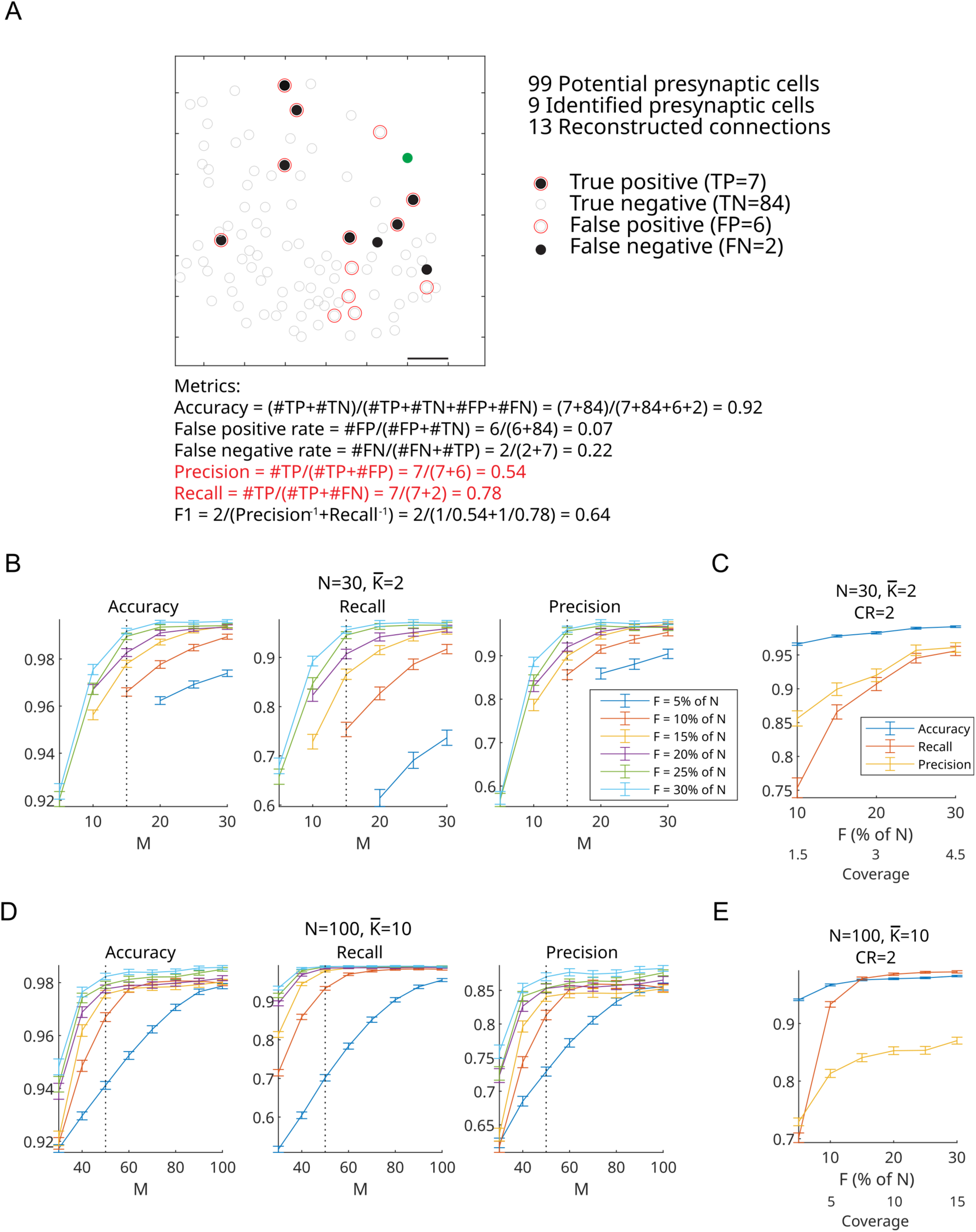
Simulation of compressive sensing network reconstruction. A) Illustration of true positive, true negative, false positive, false negative and of the metrics for accessing the performance of reconstruction in an example FOV (corresponding to Figure 4B and 5B). Filled green dots represent the patched cell, other dots represent investigated potential presynaptic cells. Filled black circles and open grey circles indicated connected and non-connected cells, identified in sequential connectivity mapping experiment. Red open circles represented reconstructed connections. B) Performances of a simulated connectivity reconstruction upon multi-cell stimulation of varying target numbers (F) in artificial networks of N=30 potential presynaptic neurons among which an average of *K̅*^)^=2 cells are connected to the monitored postsynaptic cell (connectivity rate 6.7 %). (see Methods for details). Precision, recall and accuracy are reported as a function of measurement number (M, i.e. stimulation patterns) and colors code for different number of cells stimulated per measurement (F). Dotted vertical lines indicate compression ratio of 2. C) Performances of reconstruction with compression ratio of 2, as a function of the number of cells stimulated per measurement (F) and coverage (M*F/N). Colors code for different metric (precision: blue; recall: red; accuracy: yellow). D,E) Similar to (B,C) but for larger (N=100) and denser networks (*K*^)^=10 connected cells, connection rate=10%).

**Figure S11:**
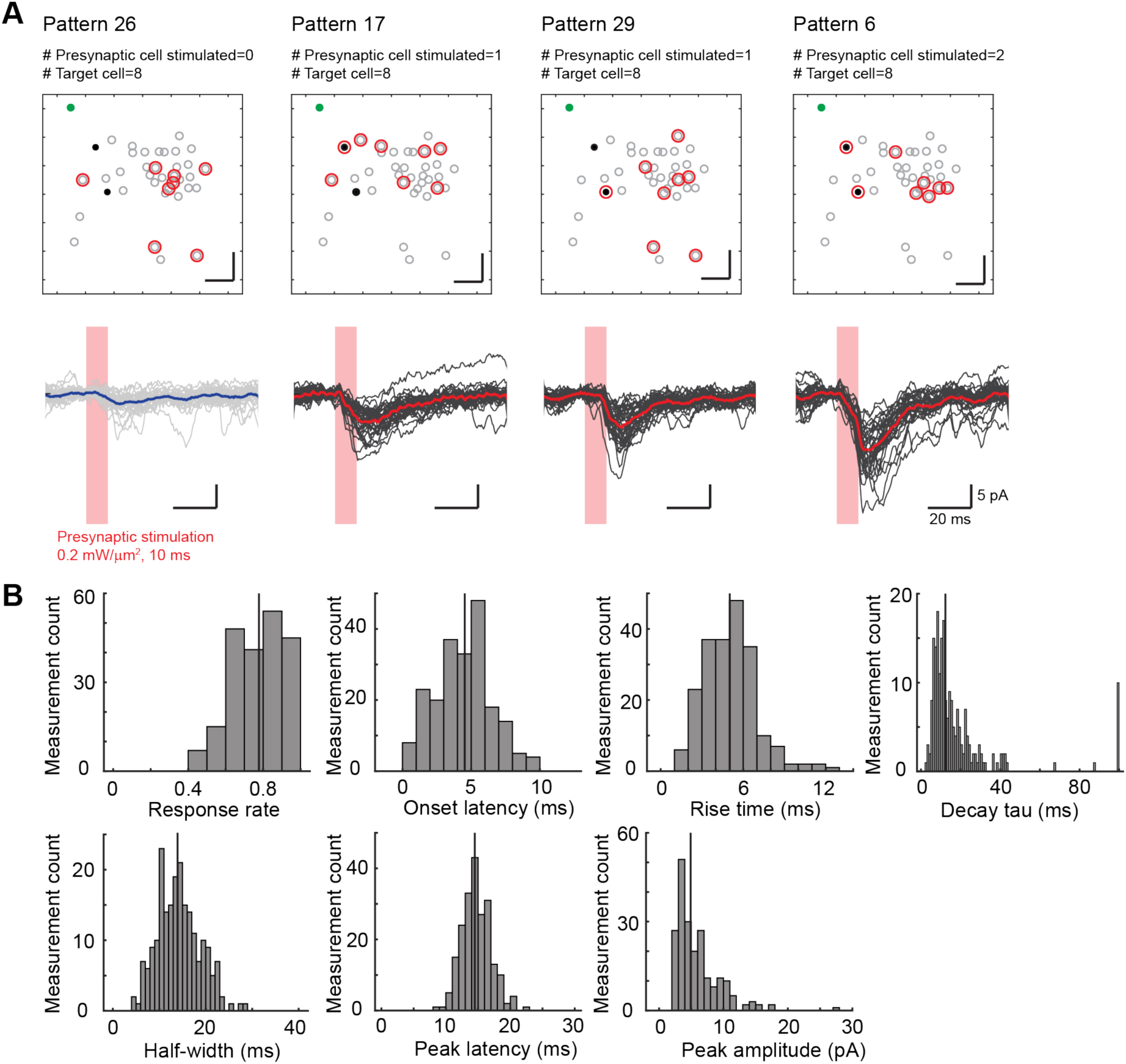
Postsynaptic activity in response to multi-cell stimulation of potential presynaptic cells. (A) Example experiment of multi-cell stimulation. Upper, maps of the coordinates of a postsynaptic cell (green circle) and 37 potential presynaptic cells (grey circles) in an experiment of multi-cell stimulation. Different sets of 8 cells were illuminated (red circles) in the same FOV as that in Figure 3A. The 2 presynaptic cells identified in sequential connectivity mapping experiment were indicated as black filled circle. The 4 represented light-patterns targeted none of (Pattern 26), each of (Patterns 17 and 29) or both of (Pattern 6) the 2 presynaptic cells. Scale bar: 50 µm. Lower, example traces of postsynaptic current in response to stimulation of the 4 light-patterns targeting different sets of 8 potential presynaptic cells. Non-responding individual traces denoted in light grey, with their average trace in blue. Responding individual traces denoted in dark grey, with their average traces in red. 32, 34, 35, 32 repetitions for stimulation of Pattern 26, 17, 29, and 6 respectively. The duration of presynaptic stimulation indicated in red shades. (B) Histograms of the properties of responding postsynaptic current upon multi-cell stimulation of potential presynaptic cells. The response rates were calculated from individual traces; other response properties were extracted from the average traces across individual postsynaptic current traces. Medians denoted as vertical lines (210 measurements of multi-cell stimulation in 12 experiments).

**Figure S12:**
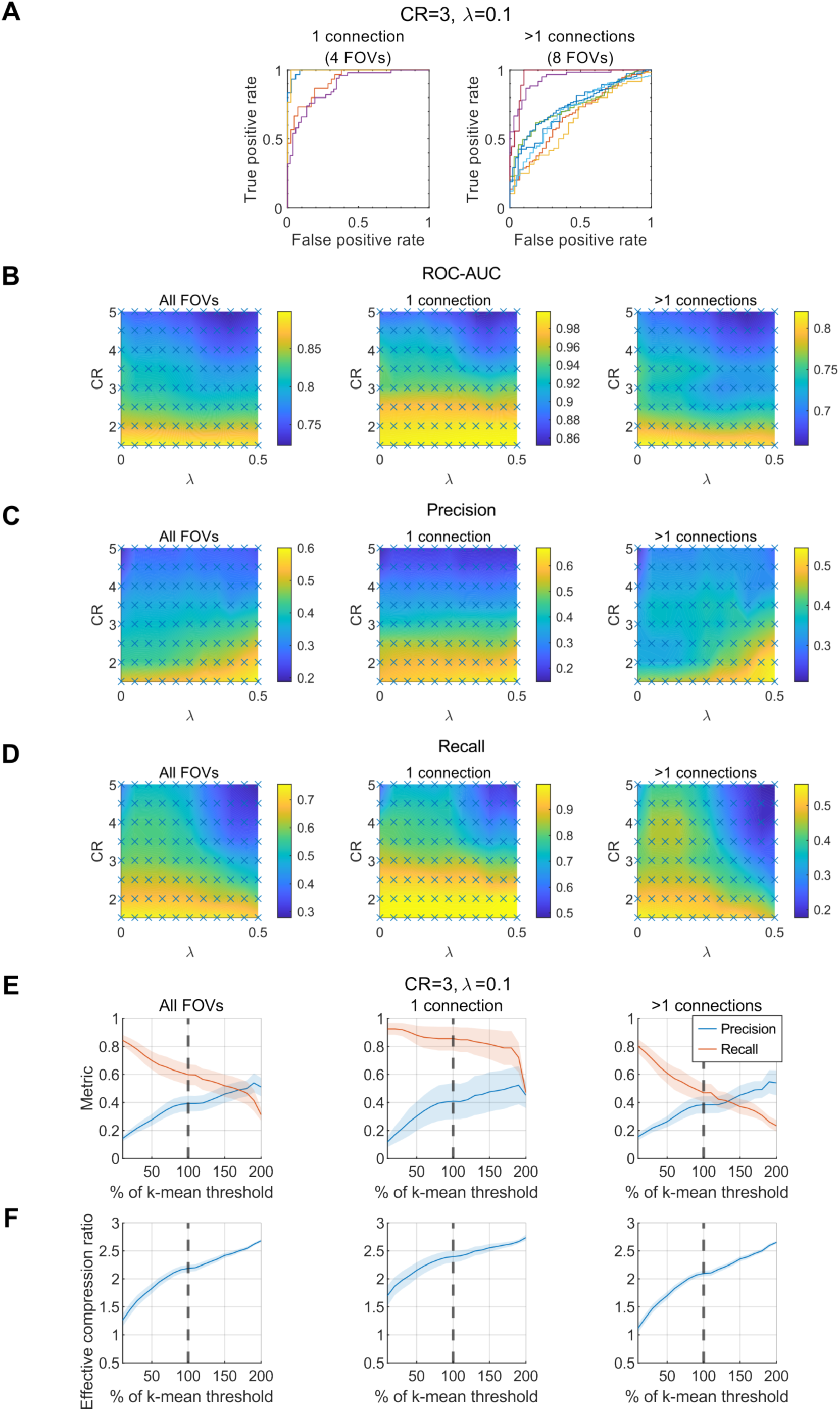
Experimental parallel connectivity mapping and reconstruction performances varying measurement numbers (m), regularization parameter λ and connection-defining threshold. (A) Receiver operating characteristic (ROC) curves visualizing the performance of CS reconstruction (see Methods), with regularization factor λ=0.1 and measurement numbers corresponding to compression ratio CR=3 for 4 FOVs of 1 connection and 8 FOVs of >1 connections. Each color represents 1 out of 12 FOVs (same datasets as reported in Figure 5C, 5D and Table 3). (B-D) Color contour map representing reconstruction performances in terms of values of the area under ROC curve-ROC-AUC (B), precision(C), and recall (D) for different values of λ and compression ratio (CR). (E) Precision and recall with different values of the connection-defining threshold used to discern connected and non-connected cells from reconstruction results (blue line in main Figure 5A, B). 100% values represent the threshold resulting from k-mean clustering of the reconstructed amplitudes (values used for results in Figure 5). The curves represent values of achievable precision (blue) and recall (red) for fixed values of CR=3 and λ=0.1, shaded area represent the standard deviation of the FOVs. By decreasing or increasing threshold, more or fewer cells are retrieved as connections, varying the number of false positive and false negative. (F) Similar to (E), with y-axis representing the effective compression ratio, which takes into account the extra number of measurements required to rule out the false positive by re-probing all the positive reconstructed connections.

**Table S3:** Comparison of presynaptic stimulation in recent studies of connectivity mapping in mouse brain by using 2P optogenetics (Hage et al., 2022, Printz et al., 2023, Izquierdo-Serra et al., 2018).

|  | Current study | Hage et al., 2022 | Printz et al., 2023 | Izquierdo-Serra et al., 2018 |
| --- | --- | --- | --- | --- |
| Investigated connectivity | Connectivity between L2/3 neurons in mouse visual cortex, biasing excitatory connections | Intralaminar and translaminar connectivity to L2/3 excitatory or inhibitory neurons in mouse visual cortex | Connectivity between neurons projecting to basolateral amygdala in mouse medial prefrontal cortex | Connectivity between L2/3 excitatory neurons in mouse visual cortex |
| Optical method | CGH and temporal focusing | spiral scanning | spiral scanning | spiral scanning |
| Opsin | Somatic ST-ChroME | Somatic or non-somatic ChrimsonR | Somatic CoChR | non-somatic C1V1 |
| Working stimulation condition | 0.15-0.3 mW/$\mu\text{m}^2$ (17-34 mW/cell), 10 ms | 35 or 85 mW/cell, 10 ms (85 mW/cell for Penk line) | 10 mW/cell after scattering, ~7 ms | 7 mW/cell, 50 ms |
| Preparation | Anesthetized mouse brain | Brain slices | Brain slices | Brain slices |
| AP properties upon single-cell stimulation | ~4 ms latency, ~0.8 ms jitter, ~1.5 count, ~80% of responding neurons, ~95% AP probability in responding neurons | ~6 ms latency, ~0.5 ms jitter, ~2.7 count, 100% probability for non-somatic ChrimsonR expression by using Penk mouse line | ~5 ms latency, ~1 ms jitter, ~1.4 count | ~30 ms latency, ~6 ms jitter, 72% of responding neurons |
| AP properties upon multi-cell stimulation | ~3 ms latency, ~0.6 ms jitter, ~1.8 count, ~94% probability ('10+1 targets') |  |  |  |
| FWHM of lateral selectivity of AP probability upon single-cell stimulation | 10 $\mu\text{m}$ | 12-23 $\mu\text{m}$ (~10 $\mu\text{m}$ for Penk:AAV, ~12 $\mu\text{m}$ for Tlx:AAV, ~12 $\mu\text{m}$ for Sst:AAV) | 24 $\mu\text{m}$ | Within 10 $\mu\text{m}$ laterally away from the soma for $\geq 0.4$ AP count and $\leq 60$ ms latency |
| FWHM of axial selectivity of AP probability upon single-cell stimulation | 56 $\mu\text{m}$ | 38-75 $\mu\text{m}$ (~50 $\mu\text{m}$ for Penk:AAV, ~40 $\mu\text{m}$ for Tlx:AAV, ~45 $\mu\text{m}$ for Sst:AAV) | 56 $\mu\text{m}$ | Within 30-40 $\mu\text{m}$ around the focal plane for $\geq 1$ AP count and $\leq 20$ ms latency |
| FWHM of lateral selectivity of AP probability upon multi-cell stimulation | 15 $\mu\text{m}$ | | | |
| FWHM of axial selectivity of AP probability upon multi-cell stimulation | 62 $\mu\text{m}$ | | | |
| Off-target activation upon single-cell stimulation | ~6% (Binary ellipsoid spiking probability function) | 3% for Sst, 21% for Tlx (Continuous spiking probability function) | ~13% (Binary ellipsoid spiking probability function) |  |
| Off-target activation upon multi-cell stimulation | ~18% |  |  |  |

